# LXR alters CD4^+^ T cell function through direct regulation of glycosphingolipid synthesis

**DOI:** 10.1101/721050

**Authors:** Kirsty E Waddington, George A Robinson, Beatriz Rubio-Cuesta, Eden Chrifi-Alaoui, Sara Andreone, Kok-Siong Poon, Iveta Ivanova, Lucia Martin-Gutierrez, Dylan M Owen, Elizabeth C Jury, Inés Pineda-Torra

**Affiliations:** Centre for Rheumatology, University College London, United Kingdom; Centre for Cardiometabolic and Vascular Science, University College London, United Kingdom; Department of Physics and Randall Division of Cell and Molecular Biophysics, King’s College London, United Kingdom

**Keywords:** liver X receptor, CD4^+^ T cell, lipid metabolism, cholesterol, glycosphingolipids

## Abstract

The liver X receptor (LXR) is a key transcriptional regulator of cholesterol, fatty acid, and phospholipid metabolism. Dynamic remodeling of immunometabolic pathways, including lipid metabolism, is a crucial step in T cell activation. Here we explored the role of LXR-regulated metabolic processes in primary human CD4^+^ T cells, and their role in controlling plasma membrane lipids (glycosphingolipids and cholesterol) which strongly influence T cell immune signaling and function. Crucially, we identified the glycosphingolipid biosynthesis enzyme glucosylceramide synthase (UGCG) as a direct transcriptional LXR target. LXR activation by agonist GW3965 or endogenous oxysterol ligands significantly altered the glycosphingolipid:cholesterol balance in the plasma membrane by increasing glycosphingolipid levels and reducing cholesterol. Consequently, LXR activation lowered plasma membrane lipid order (stability), and an LXR antagonist could block this effect. LXR stimulation also reduced lipid order at the immune synapse and accelerated activation of proximal T cell signaling molecules. Ultimately, LXR activation dampened pro-inflammatory T cell function. Finally, compared to responder T cells, regulatory T cells had a distinct pattern of LXR-target gene expression corresponding to reduced lipid order. This suggests LXR-driven lipid metabolism could contribute to functional specialization of these T cell subsets. Overall, we report a novel mode of action for LXR in T cells involving the regulation of glycosphingolipid and cholesterol metabolism, and demonstrate its relevance in modulating T cell function.

## Introduction

CD4^+^T cells (also known as T helper cells) shape the immune response by releasing cytokines with both pro-inflammatory and immunomodulatory effects. A number of factors govern the precise balance of pro- and anti-inflammatory mediators produced, including antigenic stimulation, cell-cell signaling and micro-environmental cues. The T cell plasma membrane facilitates these processes, providing a flexible interface between the cell and its microenvironment, where membrane receptors integrate internal and external signals to generate functional outcomes. Lipids are a key component of the plasma membrane and contribute to its biophysical properties and protein receptor compartmentalization. Cholesterol and glycosphingolipids are particularly enriched, forming signaling platforms known as lipid rafts which play a critical role in T cell antigen receptor (TCR) signaling and T cell function^1–3^. Cholesterol maintains lipid raft structure, inhibits spontaneous TCR activation and promotes TCR clustering^4,5^. In addition, cholesterol has been shown to regulate T cell proliferation^6,7^, cytokine production and differentiation^8^. Similarly, glycosphingolipids influence TCR-mediated signaling, responsiveness to cytokine stimulation and T_H_17 cell differentiation^9,10^. Importantly, abnormal T cell plasma membrane lipids has been linked to pathogenic T cell function and are attractive targets for immunotherapy in autoimmunity, viral infection and cancer^2,5,11,12^.

Our previous work linked pathogenic elevation of CD4^+^ T cell glycosphingolipid expression in systemic lupus erythematosus (SLE) to liver X receptor (LXR) expression^9^. LXRα (*NR1H3*) and LXRβ (*NR1H2*) are transcription factors activated by oxidized derivatives of cholesterol (oxysterols)^13^ and intermediates of cholesterol biosynthesis^14^to regulate gene expression. The majority of LXR target genes are involved in the metabolism of lipid metabolic processes, including cholesterol efflux and uptake, fatty acid biosynthesis, and phospholipid remodeling^15^. However, it is not known whether LXR regulates glycosphingolipid metabolism or T cell lipid rafts. This prompted us to further explore the relationship between LXRs, glycosphingolipid metabolism and plasma membrane lipid composition.

Here we demonstrate a new role for LXR in human CD4^+^ T cells that involves modulation of the human T cell transcriptome and lipidome. We show how LXR activation modulates glycosphingolipid and cholesterol homeostasis and define a mechanism for LXR-mediated effects on T cell function via regulation of plasma membrane lipid composition. Finally, we show that regulatory T cells (Tregs) have a distinct plasma membrane lipid profile that corresponds to differential expression of LXR target genes. We propose that regulation of membrane lipids by LXR could contribute to the specialized regulatory functions of this T cell subset.

## Results

### LXR transcriptionally regulates lipid metabolic pathways in human CD4^+^ T cells

To define the transcriptional effects of LXR activation in human CD4^+^ T cells, primary cells were exposed to the specific LXR agonist GW3965 (GW)^16^. Sixty-five LXR-responsive genes were identified (Fig. 1a-b, Dataset S1), and GW-treated samples were clearly distinguishable from their controls by principal component analysis (PCA) (Fig. S1a). The majority of differentially expressed genes (DEGs) were upregulated (53 out of 65), a subset of which demonstrated a very strong ligand response. These included well-characterised LXR target genes (*ABCG1, ABCA1, APOC1, SCD* and *SREBF1*^17^) and the recently identified oligodendrocyte maturation-associated long intervening non-coding RNA (*OLMALINC*)^18^ (Fig. 1c, Fig. S1b). The significantly enriched pathways were hierarchically clustered into functionally related groups (Fig. 1d). Strikingly, all 15 clusters enriched for LXR-upregulated genes were related to metabolism, the most significant of which was ‘cholesterol metabolic process’. Only 12 genes were significantly downregulated by GW (Fig. 1a-b, Fig. S1b), and these were most strongly associated with the ‘regulation of inflammatory responses’ (Fig. 1e).

**Figure 1.**
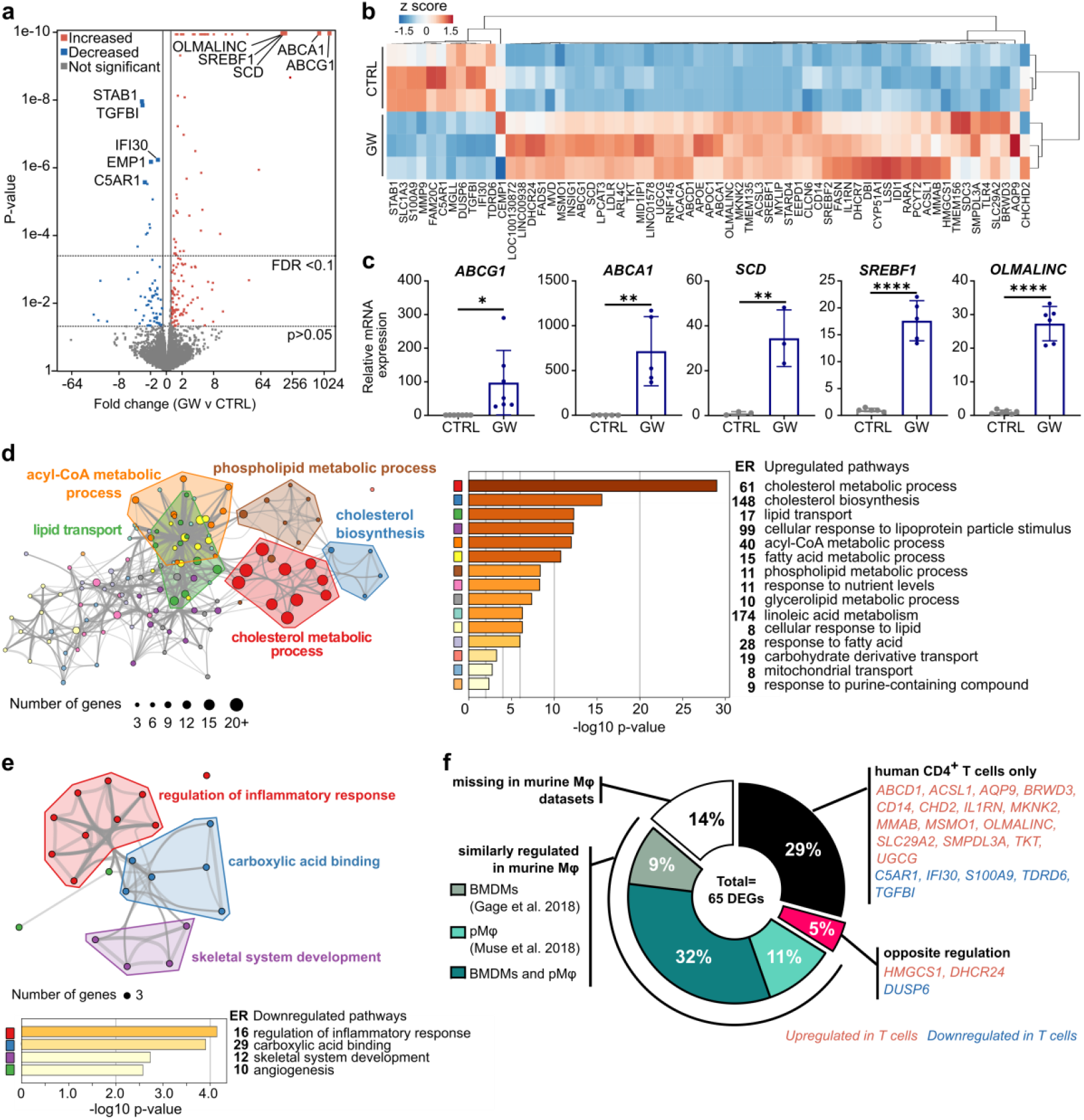
LXR regulates lipid metabolism in human CD4^+^ T cells. Primary human CD4^+^ T cells (n=3) were cultured with or without LXR agonist (GW3965, GW) for 24 hours and gene expression was assessed by RNA-Seq. **(a)** Volcano plot showing fold changes and p-values. Coloured points represent significantly regulated genes (p < 0.05). **(b)** Clustered heatmap of normalised gene counts of all LXR-regulated genes with FDR corrected p < 0.1. **(c)** Regulation of a selection of genes was confirmed by qPCR in an independent set of donors (n=3-6). Bars represent mean ± SD. Unpaired two-tailed t-test; *p < 0.05, **p < 0.01, ****p < 0.0001. **(d-e)** Network diagrams illustrate pathways significantly enriched for up- **(d)** or down- **(e)** regulated genes. Each node represents a significantly enriched term, with node size proportional to the number of contributing genes. Similar terms with a high degree of redundancy were clustered, as depicted. Bar charts plot cluster significance, and show enrichment ratios (ER). **(f)** Pie chart showing the proportion of genes regulated by GW in human T cells that are also regulated in murine bone marrow derived macrophages (BMDMs)^21^ or peritoneal macrophages (pMφ)^22^.

LXRs can act in a subtype specific manner, and the relative expression of LXRα and β differs between monocytes/macrophages and T cells^7,19^ (Fig. S1c-d). Another striking difference is that in monocytes/macrophages LXRα regulates its own expression via an autoregulatory loop^20^ which does not occur in T cells (Fig. S1e). These differences likely lead to cell-type specific responses to LXR activation. To identify potential T cell specific LXR targets we cross-referenced our list of DEGs with two publically available RNA-sequencing datasets from murine macrophages (mMφ) treated with GW^21,22^. Of the DEGs identified in T cells, 52% were similarly regulated in mMφ, and remarkably, 29% were uniquely regulated in the T cell dataset (Fig. 1f). Some of these genes are known to be differentially regulated between mice and humans/primates but, to our knowledge, a subset have not previously been associated with LXR activation (*BRWD3, CHD2, MKNK2, SLC29A2, TDRD6, TKT* and *UGCG*) (Table S1). Overall, genes involved in lipid metabolic pathways were upregulated in both cell types, but there were no shared pathways amongst the downregulated genes, which tended to be involved in the regulation of immunity and inflammation Fig. S1f). This supports that the immunomodulatory effects of LXR activation vary between cell-types and species^23^.

Thus, we have identified genes responsive to LXR activation in human CD4^+^ T cells, most markedly the upregulation of genes involved in lipid metabolic processes, and highlighted a subset of genes that may represent human or T cell specific targets.

### LXR controls transcriptional regulation of glycosphingolipid biosynthesis enzyme UGCG

Since LXR activation predominantly regulated genes involved in lipid metabolism, the impact on T cell lipid content was assessed using shotgun lipidomics. Although total intracellular lipid levels were not affected by LXR activation (Fig. 2a), 15% of the detected lipid subspecies were significantly regulated (54 out of 366, Fig. 2b). Notably, a large proportion of triacylglycerols (TAG) and hexosylceramides (HexCer) were induced by LXR activation, and overall quantities of TAG and HexCer were elevated (Fig. 2c-e, Table 1).

**Table 1.** CD3^+^ T cells were cultured ± GW3965 (GW) for 36 hours (n=4). Lipid concentrations are given in pmol and expressed as a percentage of the total lipid content (%). Samples were compared by a paired two-tailed t-test; p < 0.05 are bold. The average co-efficient of variance for each variable was used to determine the appropriate number of significant figures to display.

| Lipid class | CTRL (pmol) | GW (pmol) | *P*-value | CTRL (%) | GW (%) | *P*-value |
| --- | --- | --- | --- | --- | --- | --- |
| | Mean ± SD | | paired | Mean ± SD | | paired |
| **Cholesterol esters** | 14 ± 5 | 9 ± 1 | 0.220 | 0.06 ± 0.02 | 0.04 ± 0.00 | 0.111 |
| **Ceramide** | 60 ± 22 | 54 ± 11 | 0.411 | 0.28 ± 0.07 | 0.25 ± 0.05 | 0.065 |
| **Cardiolipin** | 251 ± 96 | 286 ± 112 | 0.138 | 1.3 ± 0.6 | 1.4 ± 0.7 | 0.289 |
| **Diacylglycerol** | 131 ± 23 | 142 ± 29 | 0.491 | 0.6 ± 0.1 | 0.7 ± 0.1 | 0.475 |
| **Hexosylceramide** | 20 ± 4 | 27 ± 2 | **0.017** | 0.10 ± 0.01 | 0.12 ± 0.01 | **0.005** |
| **Lysophospholipid** | 14 ± 1 | 13 ± 3 | 0.370 | 0.07 ± 0.01 | 0.06 ± 0.02 | **0.032** |
| **Phosphatidate** | 10 ± 1 | 13 ± 4 | 0.231 | 0.05 ± 0.01 | 0.06 ± 0.02 | 0.324 |
| **Phosphatidylcholine** | 7023 ± 1329 | 6980 ± 1764 | 0.958 | 33 ± 3 | 32 ± 6 | 0.478 |
| **Phosphatidylcholine-ether** | 346 ± 57 | 336 ± 74 | 0.788 | 1.6 ± 0.2 | 1.5 ± 0.2 | 0.184 |
| **Phosphatidylethanolamine** | 2212 ± 509 | 2352 ± 636 | 0.538 | 10 ± 2 | 11 ± 2 | 0.377 |
| **Phosphatidylethanolamine-ether** | 2350 ± 355 | 2548 ± 418 | 0.215 | 11 ± 1 | 12 ± 1 | 0.141 |
| **Phosphatidylglycerol** | 60 ± 14 | 66 ± 15 | 0.501 | 0.28 ± 0.05 | 0.30 ± 0.05 | 0.374 |
| **Phosphatidylinositol** | 3509 ± 694 | 3664 ± 626 | 0.307 | 17 ± 3 | 17 ± 4 | 0.638 |
| **Phosphatidylserine** | 3767 ± 668 | 3820 ± 663 | 0.886 | 18 ± 3 | 18 ± 5 | 0.998 |
| **Sphingomyelin** | 1249 ± 184 | 1333 ± 106 | 0.220 | 5.9 ± 0.4 | 6.2 ± 0.4 | 0.553 |
| **Triacylglycerol** | 38 ± 13 | 113 ± 46 | **0.032** | 0.18 ± 0.05 | 0.51 ± 0.17 | **0.013** |

**Figure 2.**
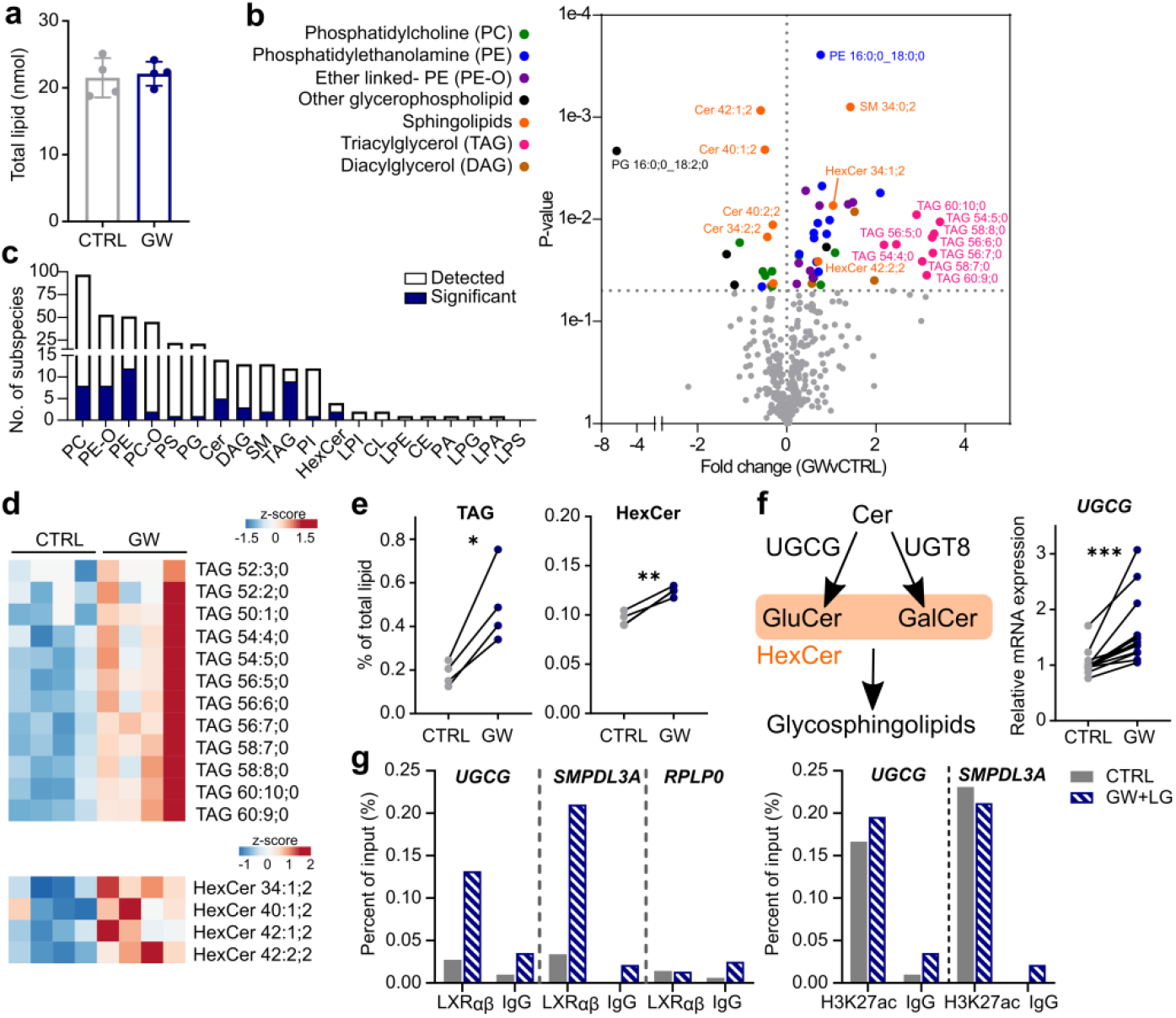
LXR activation regulates the transcription of glucosylceramide synthase (*UGCG*) **(a-e)** Primary human T cells (n=4) were treated ± GW (2 μM) for 36 hours and total cellular lipid content analysed by shotgun lipidomics. **(a)** Total lipids (normalized to cell numbers) were unchanged (mean ± SD). **(b)** Volcano plot represents significant changes in the expression of lipid sub-species, colour coded by broader lipid class (p < 0.05). **(c)** Bars show the number of subspecies detected for each lipid type. The filled area represents the proportion of subspecies significantly altered by GW treatment. **(d)** Unclustered heatmaps represent levels of individual sub-species. **(e)** Dot plots show overall change in triacylglycerol (TAG) and hexosylceramide (HexCer) levels. **(f)** Schematic illustrating the role of UGCG in the conversion of ceramide to HexCer, and upregulation of UGCG mRNA expression in CD4^+^ T cells after 24 hours GW treatment (n=13). **(g)** Cells were treated with LXR (GW, 1 μM) and RXR (LG100268; LG, 100 nM) ligands for 2 hours. LXR occupancy and H3K27 acetylation at the putative DR4 motif at *UGCG* compared to IgG control, positive control (*SMPDL3A*) and negative control (*RPLP0*) sequences. Representative of three independent experiments. **(a-f)** Two-tailed t-tests: *p < 0.05, **p < 0.01, ***p < 0.001. Abbreviations Cer - ceramide, PE - phosphatidylethanolamine, DAG - diacylglycerol, SM - sphingomyelin, PE-O - phosphatidylethanolamine-ether, PI - phosphotidylinositol, PC - phosphatidylcholine, PS - phosphatidylserine, PC-O - phosphatidylcholine-ether, LPI - lyso-phosphotidylinositol, LPE - lyso-phosphatidylethanolamine, CE - cholesterol esters, PA - phosphatidate, CL - cardiolipin, LPG - lyso-phosphatidylglycerol, LPA - lyso-phosphatidate, LPS - lyso-phosphatidylserine.

LXR regulated many enzymes involved in fatty acid metabolic processes including synthesis, desaturation and elongation (Fig 1d, Dataset S1). There were no changes in total levels of saturated, monounsaturated and polyunsaturated lipids. However, amongst PUFAs there was an increase in degree of unsaturation, which is associated with membrane disorder^24,25^ (Fig. S2a). Further examination at the lipid class level revealed significant increases in saturated and monounsaturated lipid species, HexCers, and TAG species with more than 4 double bonds (Fig. 2c and S2b).

This is the first report linking LXR activation to HexCer. We observed GW also reduced levels of several ceramides (Fig. 2c), suggesting an accelerated conversion of ceramide to HexCer - a reaction catalysed by glycosphingolipid biosynthesis enzymes UDP-glucosylceramide synthase (UGCG) or UDP-glycosyltransferase 8 (UGT8) (Fig. 2f). In support of this, UGCG mRNA expression was upregulated by LXR activation (Fig. 2f), whereas UGT8 was absent in CD4^+^ T cells (Fig. S2c). GW treatment also enhanced UGCG expression in other immune cell types, including peripheral blood mononuclear cells (PBMCs), CD14^+^ monocytes and CD19^+^ B cells (Fig. S2d). However, in monocyte-derived macrophages and THP-1 macrophages, UGCG was only modestly increased (<1.5-fold change, Fig. S2d-e). This may explain why UGCG has not been identified as an LXR target gene in previous RNA-seq and ChIP-seq experiments using macrophages ^22,26^, in which most of LXR biology has been reported to date. The increase in UGCG expression was not a GW-specific effect, as UGCG mRNA was also upregulated in response to stimulation with the endogenous LXR activators 24S, 25-epoxycholesterol (24S,25-EC) and 24S-hydroxycholesterol (24S-OHC), albeit with an altered kinetic (Fig. S2f).

To determine whether LXR regulates UGCG expression by directly binding to the UGCG locus, we screened for potential LXR response element (LXRE) sequences *in silico*. A putative DR4 sequence was identified upstream of the UGCG gene that coincided with an LXR-binding peak in HT29 cells treated with GW^27^(Fig. S2g). ChIP-qPCR experiments demonstrated enrichment in LXR occupancy at this site, which increased with ligand activation (Fig. 2g). Likewise, acetylation of histone H3K27 was enriched at this region compared to the IgG negative control, suggesting this site falls in an active transcriptional enhancer. The observed LXR occupancy at the UGCG gene followed a similar pattern to that of a reported LXRE within SMPDL3A^28^(Fig. S2g).

### LXR regulates the T cell plasma membrane lipid raft profile

UGCG is the rate-limiting enzyme for the biosynthesis of glycosphingolipids, important components of plasma membrane lipid rafts. Indeed, LXR activation consistently upregulated T cell glycosphingolipid expression measured using cholera-toxin B (CTB)(Fig. 3a), a well-established surrogate glycosphingolipid marker^9^. Specific pharmacological inhibition of UGCG activity blocked the induction of glycosphingolipids by GW, suggesting this was UGCG-dependent (Fig. S3h). The increase in glycosphingolipid levels was accompanied by significant downregulation of membrane cholesterol (Fig. 3b), likely due to the strong induction of cholesterol efflux transporters ABCA1 and ABCG1 (Fig. 1c, S3i). As expected, UGCG inhibition had no effect on the reduction of cholesterol (Fig. S3h). Overall, LXR activation significantly increased the ratio of glycosphingolipids to cholesterol (Fig. 3c).

**Figure 3.**
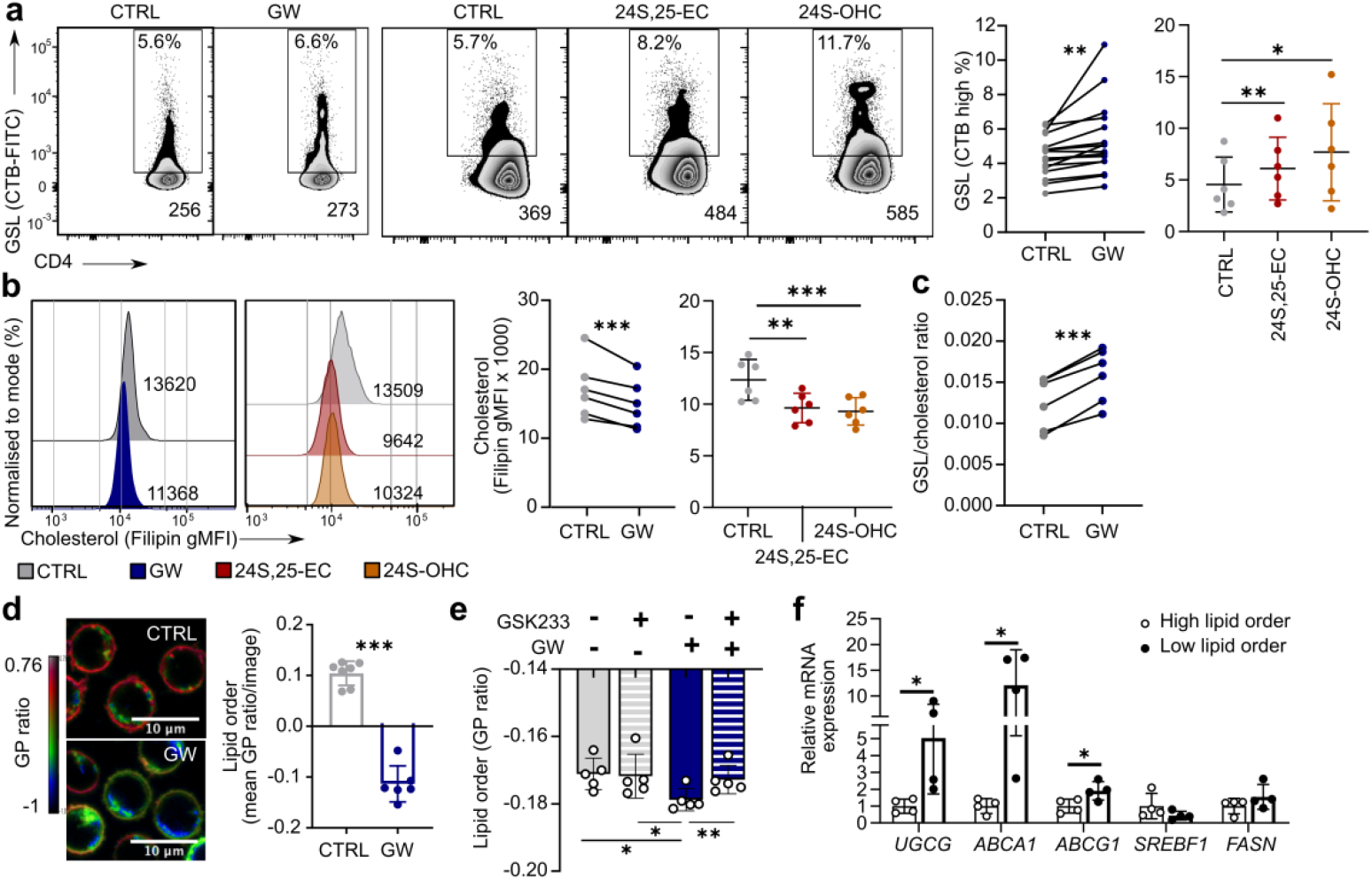
Plasma membrane lipid order is reduced by LXR activation. **(a-c)** Cells were cultured ± LXR ligands for 24 hours (GW) or 72 hours (24S,25-EC and 24S-OHC) and CD4^+^ T cell plasma membrane lipid expression was examined in >4 independent experiments. **(a)** Representative flow cytometry plots show the percentage of cells highly expressing cholera toxin B (CTB) and CTB gMFI as surrogate markers for glycosphingolipids, as in^9^. Cumulative data shows change in percentage of cells highly expressing CTB. **(b)** Representative histogram of filipin staining for cholesterol, and cumulative data showing change in gMFI. **(c)** Cumulative data showing GSL/cholesterol ratio as CTB/filipin (n=6). **(d-f)** T cell membrane lipid order was measured using di-4-ANEPPDHQ. **(d)** Representative confocal microscopy image and a histogram of average generalised polarisation (GP) ratio per image analysed are shown (n=1 donor). **(e)** Cumulative data from three experiments showing lipid order measured by flow cytometry. Cells were treated with an LXR agonist (GW) or antagonist (GSK233)(n=5) for 24 hours. **(f)** di-4**-**ANEPPDHQ-stained CD4^+^ T cells (n=4) were sorted into high or low membrane order by FACS, and gene expression was compared by qPCR. Bars show mean ± SD. **(a-f)** Two-tailed t-tests or one-way ANOVA with Tukey’s posthoc test; *p < 0.05, **p < 0.01, ***p < 0.001.

The relative abundance and arrangement of lipids in the plasma membrane dictates its ‘lipid order’, an important determinant of signalling protein localisation during immune synapse formation^29^. Cholesterol levels positively correlate with T-cell plasma membrane lipid order, whereas glycosphingolipid levels have a negative correlation^30^. LXR lowers cholesterol and raises glycosphingolipids, resulting in the significant reduction of membrane lipid order by GW/ oxysterol activated LXR (Fig. 3d-e, S3j-k). The specific LXR antagonist GSK233 was able to block the reduction of lipid order by GW, but only partially reversed the effect of 24S-OHC, in line with the known LXR-independent actions of oxysterols (Fig. 3e, S3k). Furthermore, LXR target genes were differentially expressed in T cells sorted based on their (high/low) plasma membrane lipid order. T cells with low membrane lipid order (low cholesterol, high glycosphingolipids) had elevated expression of *ABCA1, ABCG1 and UGCG* compared to T cells with high membrane lipid order (high cholesterol, low glycosphingolipids) (Fig. 3f). This suggests LXR ligand induced cholesterol efflux (*ABCA1/G1*) and glycosphingolipid biosynthesis (*UGCG*) contribute to the generation of low membrane lipid order. In contrast, there was no difference in the expression of genes controlling fatty acid synthesis (*SREBP1c, FASN*) in cells with different membrane order (Fig. 3f).

Overall, these data suggest that LXR transcriptionally upregulates *de novo* glycosphingolipid synthesis in human T cells, thereby contributing to the remodelling of plasma membrane lipid composition in response to LXR activation.

### LXR activity modulates lipid metabolism and effector function of activated T cells

Next, we explored the effect of LXR on primary human T cell activation. Over 3000 genes were significantly regulated by TCR activation, although most of these were regulated irrespective of LXR activation with GW (Fig. 4a). Interestingly, LXRβ expression was slightly increased by TCR stimulation, while LXRα expression remained low (Fig. S3a). Overall, 113 genes were regulated by the presence or absence of GW in activated T cells (Fig. 4b, S3b, Dataset S2). TCR/LXR co-stimulation upregulated genes involved in lipid metabolic processes, and downregulated genes associated with immune system processes including chemokine production and chemotaxis (Fig. S3c). When these genes were clustered based on their expression in both activated and resting cells, four major patterns of gene expression were identified (Fig. 4c, Table S2). Many genes upregulated by GW in resting cells were upregulated to an equal or greater extent in GW/TCR co-activated cells (clusters A and C, Fig. 4c). These clusters were enriched for genes involved in lipid and cholesterol metabolic processes, including canonical LXR target genes *ABCA1* and *SREBF1* and the newly identified LXR-target gene *UGCG*. This corresponded with changes in global plasma membrane lipid composition, namely increased glycosphingolipids but reduced cholesterol in response to LXR/TCR co-stimulation compared with TCR stimulation alone (Fig. 4d-f). Therefore, LXR activation continues to modulate plasma membrane composition throughout the course of T cell activation.

**Figure 4.**
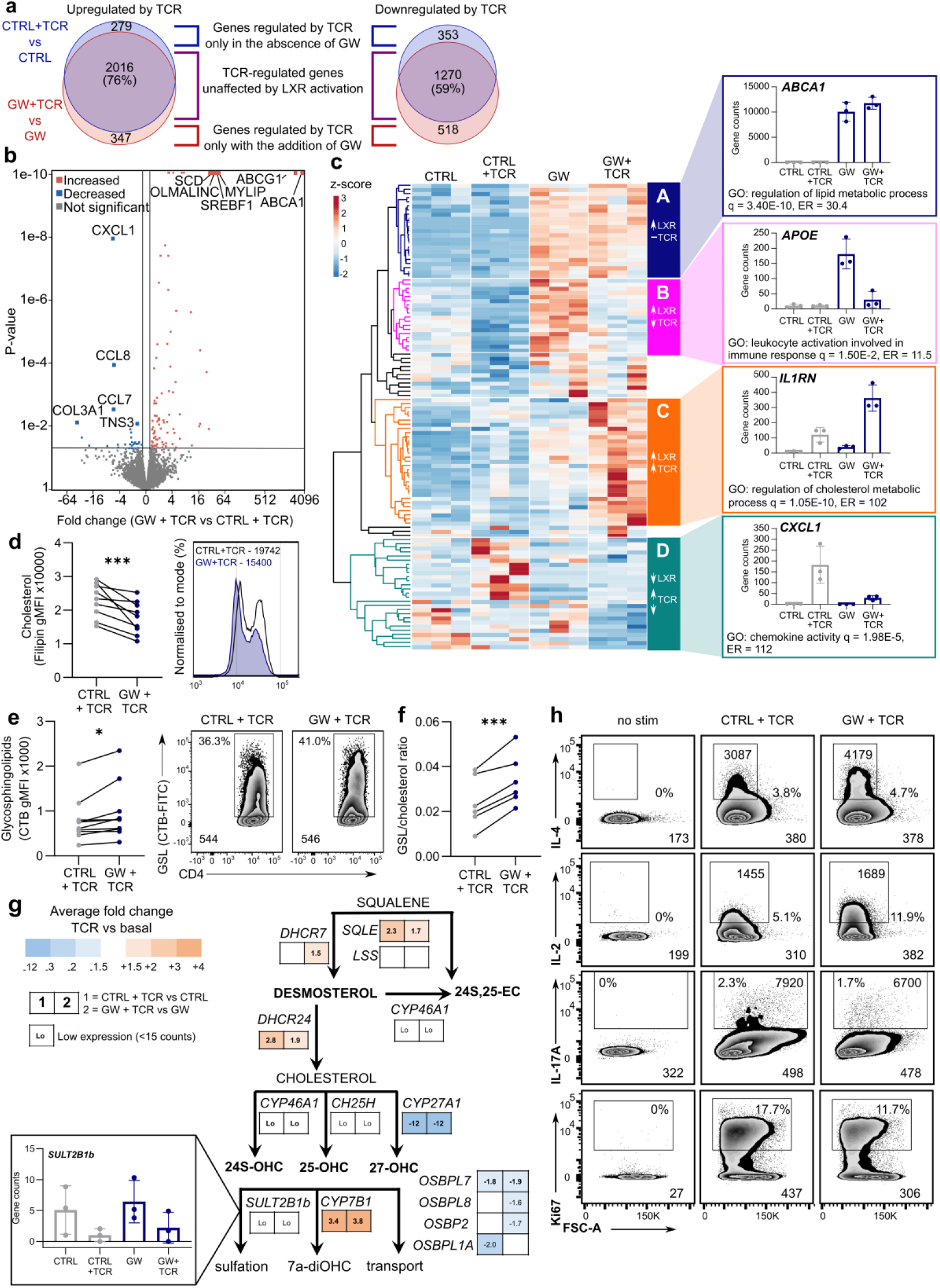
LXR activation modulates T cell immune function. **(a-c)** RNA-seq was performed on CD4^+^ T cells incubated ± GW for 6 hours, before stimulation with anti-CD3/CD28 (TCR) ± GW (n=3). **(a)** Venn diagrams compare the number of genes up or down regulated by TCR stimulation in the presence (red) or absence (blue) of GW. **(b)** Volcano plot of genes differentially expressed between GW+TCR and CTRL+TCR. **(c)** Normalized RNA-Seq gene counts of differentially expressed genes were compared to resting T cells. Four patterns of gene expression were identified by hierarchical clustering (clusters A-D). One gene from each cluster is shown as an example (mean ± SD) and the most significantly enriched gene ontology (GO) term is given. **(d-f)** Representative flow cytometry plots and cumulative data from 4 independent experiments (n=6–9) show the effect of GW on the plasma membrane cholesterol **(d)** and glycosphingolipid **(e)** content of activated CD4^+^ T cells and the ratio of GSLs to cholesterol **(f). (g)** Schematic illustrating regulation of oxysterol metabolism genes upon TCR activation. Colour coding and numbers illustrate fold change (CTRL+ TCR vs CTRL and GW+TCR vs GW). White squares represent no significant change. Bar chart plots normalized RNA-seq gene counts for *SULT2B1* (mean ±SD). **(h)** Cytokine production was analysed by intracellular flow cytometry after 72 hours anti-CD3/CD28 and 5 hours stimulation with PMA and ionomycin. Ki67 was used as a marker of proliferation. Representative flow cytometry plots are labelled with percentage of positive cells and gMFI of both the cytokine-producing population and total T cells. Two-tailed t-tests; *p < 0.05, ***p < 0.001.

In contrast, GW/TCR co-stimulation reduced the induction of a subset of genes involved in leukocyte activation (cluster B, Fig. 4c). Interestingly, other genes were only activated (cluster C) or repressed (cluster D) by LXR activation in the context of TCR stimulation (Fig. 4c). Therefore, bidirectional crosstalk between LXR and TCR stimulation modulates transcription in a gene-specific manner. Likely, more subtle differences did not reach statistical significance due to the heterogeneous response to stimulation between the healthy donors (Fig. S3d).

In murine T cells, TCR stimulation was previously reported to repress LXR transcriptional activity, by reducing the availability of endogenous LXR ligands due to their modification by the sulfotransferase SULT2B1^7^. However, in the present study we observed very low levels of *SULT2B1* in human CD4^+^ T cells (<11 gene counts), and *SULT2B1* was not regulated by TCR activation (Fig. 3g). We considered that oxysterol levels could be controlled by an alternative mechanism, for example increased efflux or metabolism. Indeed, TCR activation downregulated the expression of oxysterol-binding proteins and oxysterol biosynthesis enzyme *CYP27A1*, and upregulated oxysterol metabolising enzyme *CYP1B1* (Figure 3g).Therefore concentrations of endogenous LXR ligands during human T cell activation are also tightly regulated, but likely through a different mechanism.

LXR and T cell co-activation had significant functional consequences including increased production of interleukin (IL)-2 and IL-4, reduced IL-17A release and inhibited proliferation compared to non-LXR-treated controls (Fig. 4h, Fig. S4a-c). No changes in T cell interferon-γ, tumour necrosis factor-α or IL-10 production were detected (Fig. S4d). Although LXR has been regulate to regulate the transcription of certain cytokines^8,31^, this was not observed here (Dataset S2). Considering the preferential regulation of lipid metabolism genes (Fig. S3c) and observed changes in plasma membrane lipid levels (Fig. 4d-f), we instead hypothesised that the effects of LXR activation on T cell function could be mediated by an altered lipid landscape.

### LXR-driven modification of plasma membrane lipid profile alters TCR signalling

T cell activation is initiated by TCR-proximal signalling at the immune synapse, leading to proliferation and cytokine production. We previously demonstrated that, compared to cells with highly ordered plasma membranes, T cells with lower membrane lipid order have reduced synapse area, transient synapse formation and a Th1 cytokine skew^30^. To examine the effect of LXR stimulation on the kinetics of lipid reorganisation during the early stages of T cell activation we used di-4-ANEPPDHQ staining and TIRF microscopy to assess the interaction between CD4^+^ T cells and antibody-coated glass coverslips (mimicking the ‘immune synapse’) (Fig. 5a, Movies S1-2). T cells pre-treated with GW had a significantly lower membrane order (generalised polarisation (GP) ratio) at the cell/coverslip interface for up to 20 minutes post-activation (Fig. 5b, Movies S1-2). However, synapse area was unaffected (Figure S4e). This was accompanied by increased levels of global tyrosine phosphorylation (Fig. 5c), increased accumulation of Lck receptor tyrosine kinase at the synapse (Fig. 5c) and a preference for Lck to accumulate at the synapse periphery (Fig. 5d), an area typically associated with active signalling^32^. Specifically, GW treatment increased phosphorylation of important proximal T cell signalling molecules CD3 and the adaptor molecule linker for activation of T cells (LAT), but not extracellular signal related kinase (Erk) or phospholipase (PL) Cγ1 (Fig. S4f).

**Figure 5.**
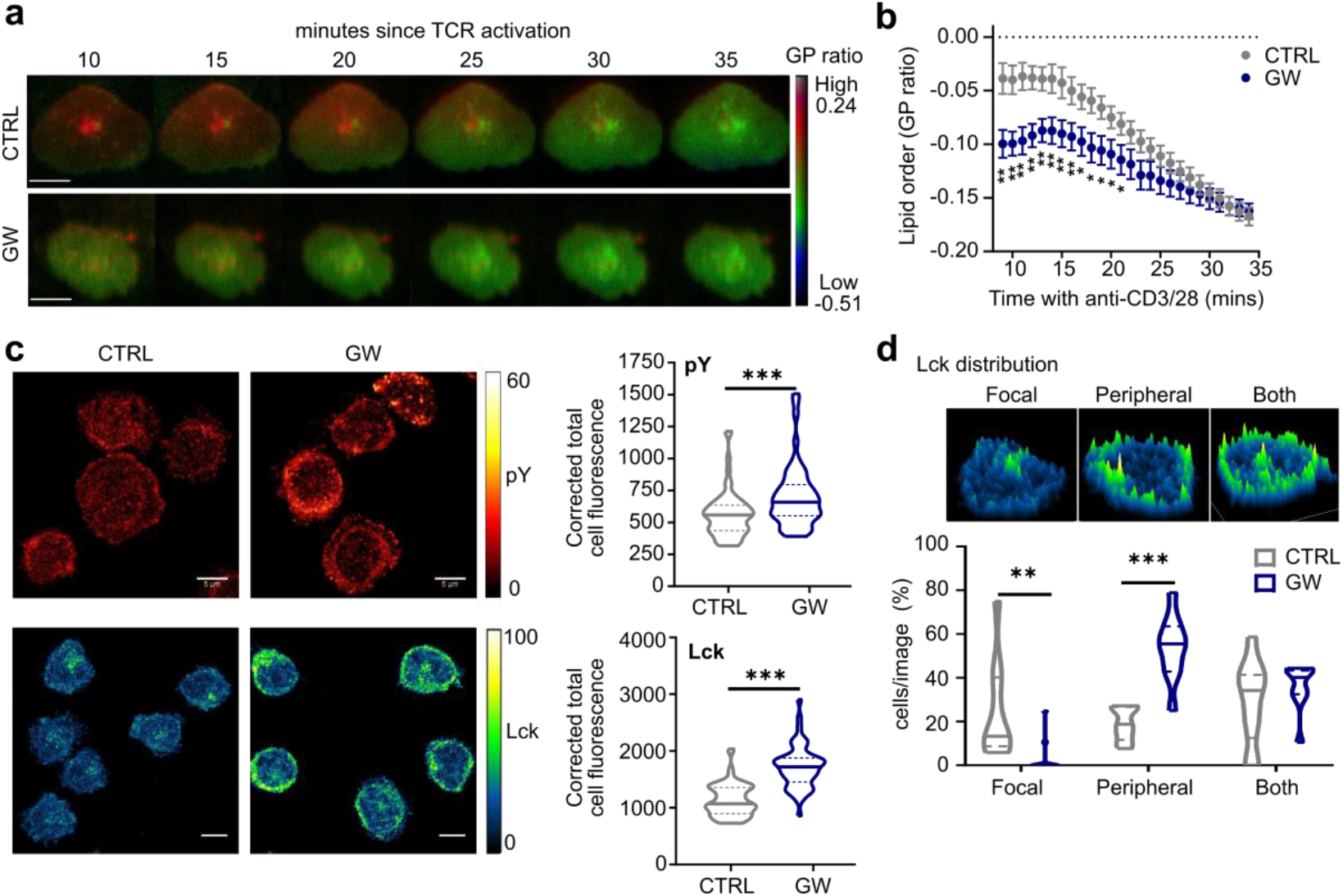
LXR activation regulates immune synapse formation and proximal TCR signalling. **(a-d)** CD4^+^ T cells were cultured ± GW before addition to chamber slides coated with anti-CD3/28 for immune synapse formation. **(a-b)** T cells were stained with di-4-ANEPPDHQ and immune synapse formation was recorded for 30 minutes using TIRF microscopy. **(a)** Representative images at 5 minute intervals, scale bar = 5 µM. **(b)** GP ratio was quantified at each minute (n= 10-12 cells/condition, mean ± SEM). **(c-d)** Immune synapses (n=2 donors) were fixed at 15 mins post activation and immunostained for Lck (CTRL=68 cells, GW=52 cells) and phosphotyrosine (pY) (CTRL=59 cells, GW=52 cells). Representative images and quantification of corrected total cell fluorescence (CTCF) **(c)** or classification of Lck distribution patterns **(d)**. Violin plots show median and quartile values. Multiple unpaired t-tests corrected for multiple comparisons **(b)** or Mann Whitney U **(c-d)**: **\***p < 0.05, **p < 0.01, ***p < 0.001. Abbreviations: Lck – lymphocyte-specific protein tyrosine kinase; pY – phosphotyrosine.

Taken together, these results suggest that plasticity in T cell function could be driven, at least in part, by altered plasma membrane lipid composition controlled by LXR activation.

### Functional T cell subsets differ in their expression of LXR-regulated genes and lipids

T cells with high and low membrane lipid order are functionally distinct ^30^. Compared to responder T cells (Tresp), regulatory T cells (Treg) (Fig. 6a) had lower membrane order increased glycosphingolipid levels and reduced membrane cholesterol (Fig. 6b-d). We hypothesised that the LXR pathway could contribute to these differences. LXRα mRNA expression was significantly lower in Tregs, although LXRβ, which is the predominant form in T cells (Fig. S1d-f), tended towards higher expression (p = 0.06)(Fig. 6e). Corresponding to the plasma membrane lipid phenotype, Treg expression of the cholesterol transporter *ABCG1* and glycosphingolipid enzyme *UGCG* were increased compared to Tresp, whereas other LXR target genes were not differentially expressed (*ABCA1, IDOL, SREBF1, FASN)* (Fig. 6e).

**Figure 6.**
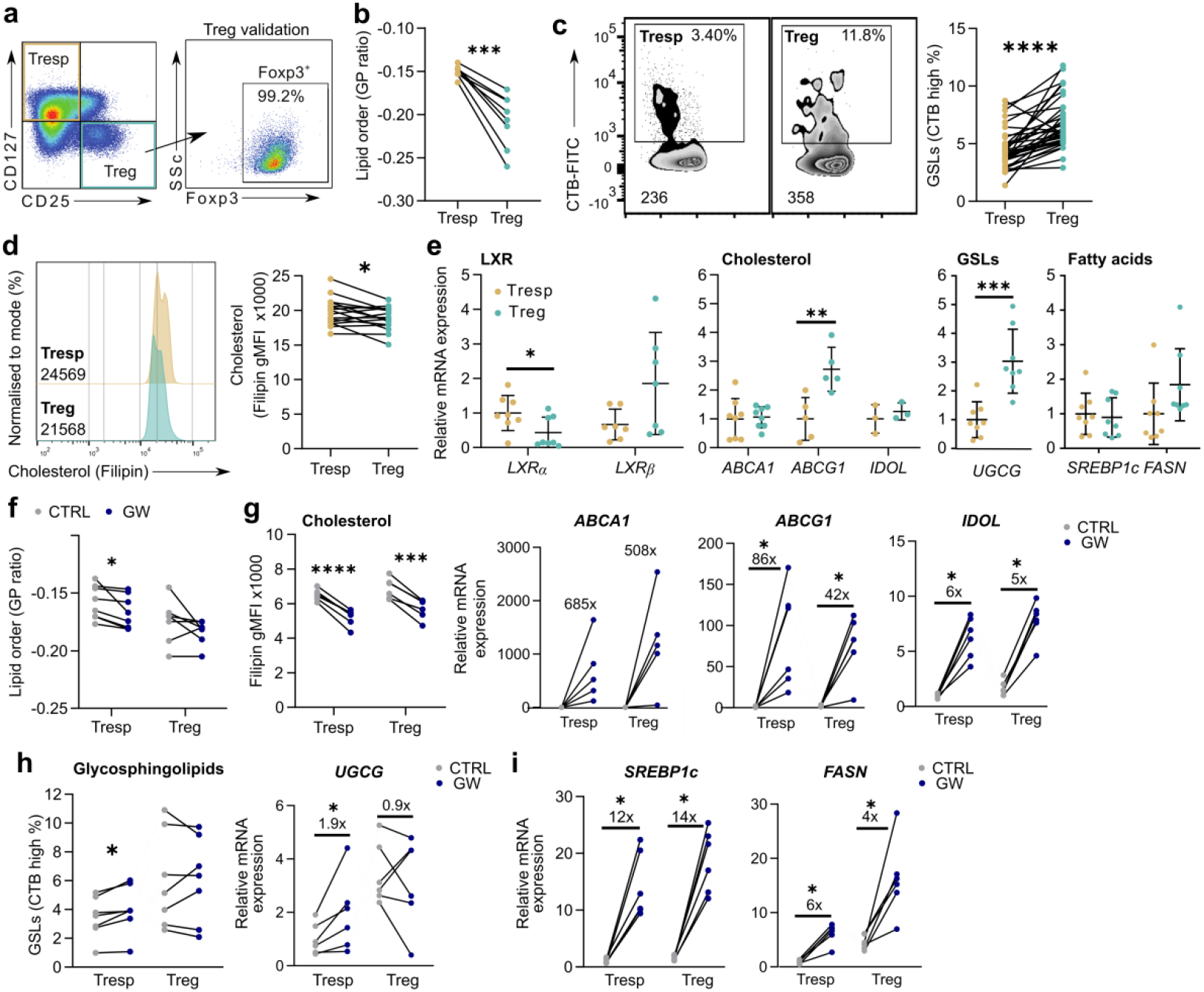
Treg and Tresponder subsets have distinct lipid metabolic phenotypes. **(a)** Responder (Tresp: CD4^+^CD25^lo^CD127^+^) and regulatory (Treg: CD4^+^CD25^+^CD127^-^) T cell subsets were defined by flow cytometry. **(b-d)** Plasma membrane lipid order (GP ratio) **(b)**, glycosphingolipid levels (GSL) **(c)**, and cholesterol content **(d)** were analysed using flow cytometry. Lines connect matched Tresp and Treg results from the same sample. **(e)** Expression of LXR and LXR-target genes that regulate cholesterol, GSL, and fatty acid levels were analysed in FACS sorted T cell subsets (n=3-8). Mean ± SD. **(f-i)** Cells were treated with GW for 24 hours. Lines connect control (CTRL) and GW treated samples from the same donor. Cumulative data from 3 independent experiments shows the change in membrane lipid order (GP ratio) **(f)**, cholesterol **(g)**, and GSL **(h)** expression. **(g-i)** Induction of LXR target genes involved in cholesterol **(g)**, GSL **(h)** and fatty acid metabolism **(i)** were analysed in FACS sorted T cell subsets (n=5-6). Gene expression is expressed relative to the average of control (CTRL) treated Tresp. The average fold change (GW vs CTRL) was calculated for each subset. Two-tailed t-tests; *p < 0.05, **p < 0.01, ***p < 0.001, ****p < 0.0001.

Interestingly, Tregs had a more variable response to LXR stimulation than Tresp in terms of reduction of membrane lipid order and induction of glycosphingolipids, although downregulation of cholesterol was consistently similar (Fig. 6f-h). Mirroring the regulation of glycosphingolipids and cholesterol, cholesterol metabolism genes (*ABCA1, ABCG1, IDOL*) were similarly induced in both subsets whereas UGCG mRNA was significantly upregulated in Tresp but not Treg, whereas (Fig. 6g-h). Fatty acid synthesis enzymes had a similar magnitude of regulation (4-fold vs 6-fold), although *FASN* levels were much higher in GW-treated Treg than Tresp (Fig. 6i).

These results demonstrate that Treg and Tresp have distinct plasma membrane lipid profiles and differences in LXR ligand responses. This suggests that variation in LXR activity could influence the functional specialisation of T cell subsets.

## Discussion

CD4^+^ T cells provide essential protection against infection and cancer, but dysregulated T cell responses contribute to the pathogenesis of many diseases. LXRS are an attractive therapeutic target in many immunometabolic diseases involving T cells^33,34^. However, the actions of LXR in lymphocytes have not yet been fully investigated, particularly in human cells. This is important since a number of differences in LXR biology have been reported between human and rodent models, including the aforementioned species-specific regulation of certain genes^18,20,28,35^. Furthermore, in stark contrast to the anti-inflammatory effects of LXR activation in murine macrophages^36–38^, LXR has repeatedly been shown to potentiate pro-inflammatory responses in human monocytes^35,39,40^.

Here, we have comprehensively assessed the action of LXR in human CD4^+^ T cells combining transcriptomic and lipidomic analyses with cell biology approaches to study the regulation of lipid metabolism and T cell function. Our findings revealed a novel regulation of glycosphingolipid biosynthesis by LXR in these cells, which may be replicated in other immune cell types. The combined effect of LXR activation on glycosphingolipid and cholesterol levels contributed to an overall reduction in plasma membrane lipid order, which modulated immune synapse formation and proximal T cell signalling in the context of TCR activation.

Whilst this work was ongoing, it was demonstrated that LXR contributes to T cell development in animal models. T cell specific deletion of LXR resulted in peripheral lymphopenia, thought to be caused by accumulation of plasma membrane cholesterol, heightened lipid-raft-associated apoptotic signalling, and subsequent enhanced negative selection^41^. This supports our findings that regulation of plasma membrane lipids by LXR is important for T cell function.

The discovery that LXR activation upregulates UGCG expression in immune cells provides a novel mode of action for LXR in the immune system. UGCG is a ubiquitously expressed and highly conserved gene. To date no post translational modifications have been identified, and transcriptional regulation appears to be the main determinant of its activity^42^. UGCG expression has been shown to be strongly upregulated by a variety of inflammatory signals^43–46^, in response to inhibition of prenylation by statin treatment^47^, and by mTORC2 during tumorigenesis^48,49^. It will be important to establish whether LXR-mediated regulation of UGCG extends to other cell-types and tissues, as this could have wide-reaching implications for the therapeutic activation of LXR in various contexts. For example, elevated expression of UGCG has repeatedly been linked to acquisition of multi-drug resistance and resistance to apoptosis in an array of cancer models^50–52^. More recently, UGCG overexpression has been shown to drive enhanced glutamine and mitochondrial metabolism in breast cancer cells^53,54^.

LXR activation can be pro- or anti-inflammatory depending on the timing of stimulation and species studied^35,39,40^. LXR activation has previously been reported to inhibit cytokine production by T cells^31,55,56^, generally attributed to repression of cytokine mRNA transcription^31,55^. We confirmed inhibition of proliferation and IL-17 production as previously observed^7,31,55,56^. However, we detected an increase in the production of both IL-2 and IL-4 and, in contrast to previous studies, did not observe inhibition of IFN-γ or TNF-α. Because the anti-inflammatory actions of LXR are context dependent^35,40,57^, it is likely that differences in the conditions for T cell stimulation or LXR activation could explain this discrepancy. In addition, age and sex can both influence LXR signalling^58,59^. Future studies could explore whether these factors are relevant in LXR-dependent regulation of T cells.

In our experimental set up, LXR activation by GW did not significantly alter the induction of cytokine mRNA expression. Instead, the most significantly regulated transcriptional pathways were related to lipid metabolism, and we observed changes in plasma membrane lipid expression early (mins) and late (72 hours) in the course of T cell activation. Thus, our findings support an important role for plasma membrane lipids in mediating the functional effects of LXR activation. This mechanism is likely to be complementary to others modes of LXR action, including the transcriptional regulation of certain cytokines^8,31^ and modulation of endoplasmic reticulum cholesterol content^7^.

Finally, we identified that LXR-regulated genes and lipids were differentially expressed in Tregs. LXR has been suggested to increase Foxp3 expression and inducible-Treg differentiation in murine cells^60^. In contrast, LXR activation was recently shown to decrease the frequency of a subset of T cells, intestinal RORγt^+^ Tregs, but this was attributed to an indirect effect on myeloid cells^61^. In any case, a potential interaction between LXR signalling, plasma membrane lipids and Tregs has not yet been explored. Murine Tregs also have low membrane order, and genetic deletion of ceramide synthesizing enzyme *smpd1* increases the frequency and suppressive capacity of Tregs^62^. This supports a relationship between ceramide metabolism (in which UGCG plays a key role), plasma membrane lipid order, and Treg function. Although plasma membrane cholesterol has been shown to play an important role in the differentiation of Tregs^63^, increasing plasma membrane cholesterol was reported to have no effect on their suppressive function^64^. In contrast, reduction of intracellular cholesterol by 25-hydroxycholesterol or statin treatment inhibited Treg proliferation and expression of the immune checkpoint receptor CTLA-4^65^. Together, this work supports the hypothesis that LXR could contribute to Treg function via modulation of plasma membrane lipid order.

In addition to the changes in cholesterol and glycosphingolipid metabolism explored here, triacylglycerol (TAG) levels were also substantially upregulated by LXR activation. Compared to conventional T cells, Tregs are lipid-enriched and have increased TAG synthesis and a greater concentration of lipid droplets which serve as a fuel source and protect against lipotoxicity^66^. Furthermore, TAG also promote IL-7 mediated memory CD8^+^ T cell survival^67^. Thus, the role of LXR-driven TAG biosynthesis in T cells also warrants further investigation, although this was beyond the scope of our current study.

In their resting state, T cells express low levels of endogenous LXR ligands^68^. In our experiments, CYP27A1 was the only oxysterol synthesising enzyme consistently expressed in these cells. However, there is evidence that certain polarisation conditions can lead to dramatic regulation of oxysterol synthesis and thus endogenous modulation of LXR signalling. For example, *in vitro* differentiated type 1 regulatory cells upregulate 25-hydroxychoelsterol to limit IL-10 production^68^. In contrast, Th17 cells upregulate an enzyme that sulfates oxysterols (SULT2B1), thereby inactivating them as LXR ligands and driving preferential activation of RORγt instead of LXR^69^. LXR also plays a unique role in a subset of IL-9 producing CD8^+^ T cells (Tc9), in which cholesterol/oxysterol are tightly supressed to prevent transrepression of the *Il9* locus by LXR^8^.

Furthermore, changes in oxysterol availability have been documented in many diseases, including accumulation in atherosclerotic plaques^70^, production in the tumour microenvironment^71^, and reduced circulating levels in multiple sclerosis^72^. Therefore, the new mechanism described here could be of therapeutic relevance to disorders characterised by defects in T cell signalling and lipid metabolism. For example, in addition to altered oxysterol levels, multiple sclerosis patients are reported to have altered LXR signalling, cholesterol levels and glycosphingolipid metabolism^73–75^. However, whether plasma membrane lipid rafts contribute to immune-cell dysfunction is multiple sclerosis is currently unknown.

In conclusion, our findings show for the first time that LXR regulates glycosphingolipid levels, which strongly impacts plasma membrane lipid composition and T cell function. This new mechanism could be of therapeutic relevance to disorders characterised by defects in T cell signalling and metabolism, including autoimmune and neurodegenerative diseases, cardiovascular disease, and cancer.

## Materials and Methods

### Antibodies and reagents

#### Western blotting

Primary antibodies were anti-LXRα (clone PPZ0412, Abcam Cat# ab41902, RRID:AB_776094), anti-LXRβ (Active Motif Cat# 61178, RRID:AB_2614980), anti-phosphotyrosine (clone 4G10, Millipore Cat# 05-321, RRID:AB_309678) anti-phospho-LAT (Y191) (Cell Signaling Technology Cat# 3584, RRID:AB_2157728), anti-phospho CD3ζ (Y142) (clone EP265(2)Y, Abcam Cat# ab68235, RRID:AB_11156649), anti-p42/p44 MAPK(Thr202/204) (Cell Signaling Technology Cat# 9101, RRID:AB_331646) and anti-HSP90 (clone H-114, Santa Cruz Biotechnology Cat# sc-7947, RRID:AB_2121235). Horseradish peroxidase conjugated secondary antibodies were goat anti-rabbit IgG (Agilent Cat# P0448, RRID:AB_2617138 or Cell Signaling Technology Cat# 7074, RRID:AB_209923) or sheep anti-mouse IgG (GE Healthcare Cat# NA931, RRID: AB_772210).

### Confocal microscopy

Primary antibodies were anti-phosphotyrosine (clone 4G10, Millipore Cat# 05-321, RRID:AB_309678) and anti-Lck (Santa Cruz Biotechnology Cat# sc-13, RRID:AB_631875). Secondary antibodies were goat anti-mouse IgG2b-AlexaFluor633 (Thermo Fisher Scientific Cat# A-21146, RRID:AB_2535782) and goat anti-rabbit IgG-AlexaFluor488 (Thermo Fisher Scientific Cat# A-11034, RRID:AB_2576217).

#### Flow cytometry

(i) Cholesterol and glycosphingolipid detection: CD4-BV711 (clone RPA-T4, BioLegend Cat# 300558, RRID:AB_2564393), CD25-BV510 (clone MA251, BD Biosciences Cat# 563351, RRID:AB_2744336), CD127-PEDazzle594 (clone A019D5, BioLegend Cat# 351336, RRID:AB_2563637), CD19-APC (clone HIB19, BioLegend Cat# 302212, RRID:AB_314242), CD14-efluor450 (clone 61D3, Thermo Fisher Scientific Cat# 48-0149-42, RRID:AB_1272050), cholera Toxin B subunit FITC conjugate (Sigma-Aldrich Cat# C1655), filipin complex from *Streptomyces filipinensis* (Sigma-Aldrich Cat# F9765, CAS: 11078-21-0). (ii) Lipid order experiments: CD4-BUV395 (clone SK3, BD Biosciences Cat# 563550, RRID:AB_2738273), CD25-APC (clone BC96, BioLegend Cat# 302610, RRID:AB_314280) and CD127-BV421 (clone A019D5, BioLegend Cat# 357424, RRID:AB_2721519), di-4-ANEPPDHQ (Thermo Fisher Scientific Cat# D36802). (iii) Intracellular staining: IFNγ-efluor450 (clone 4S.B3, Thermo Fisher Scientific Cat# 48-7319-42, RRID:AB_2043866), IL-4-APC (clone MP4-25D2, BioLegend Cat# 500812, RRID:AB_315131), IL-17A-AlexaFluor488 (clone eBio64Dec17, Thermo Fisher Scientific Cat# 53-7179-42, RRID:AB_10548943), IL-10-PE (clone JES3-19F1, BD Biosciences Cat# 554706, RRID:AB_395521), TNFα-BV421 (clone Mab11, BioLegend Cat# 502932, RRID:AB_10960738) and IL-2-FITC (clone 5344.111, BD Biosciences Cat# 340448, RRID:AB_400424). (iv) Proliferation: PE mouse anti-human Ki67 set (BD Biosciences Cat# 556027, RRID:AB_2266296) or Cell Trace Violet reagent (Invitrogen).

#### Fluorescence activated cell sorting (FACS)

CD14-v450 (clone MφP9, BD Biosciences Cat# 560349, RRID:AB_1645559), CD8a-FITC (clone RPA-T8, BioLegend Cat# 301006, RRID:AB_314124), CD19-APC-Cy7 (clone SJ25C1, BD Biosciences Cat# 557791, RRID:AB_396873), CD4-BV605 (clone OKT4, BioLegend Cat# 317438, RRID:AB_11218995).

### Human samples

50 mL of peripheral blood was collected from healthy controls (HCs). Men and women aged 18-60 were recruited. Exclusion criteria included current illness/infection, statin treatment, pregnancy, breast-feeding, or vaccination within the past 3 months. For RNA-sequencing and lipidomic analysis of T cells from HCs (Fig. 1) blood leukocyte cones were purchased from NHS Blood and Transplant. Peripheral blood mononuclear cells (PBMCs) were separated on Ficoll-Paque PLUS (GE Healthcare) using SepMate tubes (StemCell Technologies). PBMCs were cryopreserved in liquid nitrogen until use. Ethical approvals for this work were obtained from the London - City & East Research Ethics Committee (reference 15-LO-2065), Yorkshire & The Humber - South Yorkshire Research Ethics Committee (reference 16/YH/0306), South Central - Hampshire B Research Ethics Committee (reference 18/SC/0323). All participants provided informed written consent.

### Cell subset purification

#### Fluorescence activated cell sorting (FACS)

CD3^+^ T cells for lipidomics analysis were sorted by FACS. Cells were washed in MACS buffer (PBS with 2% FBS (Labtech) and 1 mM EDTA (Sigma)) before staining with antibodies against surface markers for 30 minutes. Sorting was performed on a BD FACSAria II.

#### Magnetic assisted cell sorting (MACS)

CD4^+^ T cells and CD19^+^ B cells were negatively isolated using magnetic bead based separation (EasySep, StemCell Technologies). CD14^+^ monocytes were positively selected (EasySep, StemCell Technologies). Sample purities were similar to those reported by the manufacturer (95.1 ± 1.3% for negative selection and 97.6 ± 0.21% for positive selection). To obtain monocyte-derived macrophages (MDMs), monocytes were plated in low-serum media (1% FBS) for 1-2 hours in 12-well Nunc-coated plates (ThermoFisher Scientific) to promote adherence, then cultured for 7 days in complete media (RPMI 1640 (Sigma) supplemented with 10% heat-inactivated fetal bovine serum (FBS) (Labtech) and 20 μg/mL gentamycin (Sigma)).

### THP-1 cell line

The human monocytic THP-1 cell line was a gift from Jenny Dunne (UCL, UK). Cells were maintained at a density of 0.25 – 1×10^6^/mL in complete media at 37°C, 5% CO_2_. To induce monocyte to macrophage differentiation cells were plated at 0.4×10^6^/mL) in complete media supplemented with 25 ng/mL phorbol 12-myrisate 13-acetate (PMA, Sigma) for 24 hours. Cells were washed and rested overnight prior to LXR activation.

### Cell culture

#### Culture with LXR ligands

PBMCs or purified T cells were cultured in in 96 well plates (1 ×10^6^ cells/well) in complete media. Cells treated with GW3965 (GW) (1 µM, Sigma-Aldrich, CAS: 405911-17-3) were compared either to vehicle (dimethylsulfoxide, Sigma Cat# D2650), with the exception of RNA-sequencing experiments and di-4-ANEPPDHQ confocal imaging (Fig. 3d, S2j) where the LXR antagonist GSK1440233 (1 µM, GSK233) was used as control. 24S-hydroxycholesterol (10 µM, Enzo Cat# BML-GR230-0001, CAS: 474-73-7) and 24S-25,epoxycholesterol (10 µM, Enzo Cat# BML-GR231-0001, CAS: 77058-74-3) were compared to vehicle (ethanol, Sigma Cat# E7023). The UGCG inhibitor N-Butyldeoxynojirimycin (NB-DNJ, Sigma Cat#B8299, CAS: 72599-27-0) was used at a final concentration of 10 µM, as in^9^. For chromatin immunoprecipitation experiments cells were also treated with RXR agonist LG100268 (LG) (100 nM, Sigma Cat# SML0279, CAS:153559-76-3). For western blotting experiments cells were treated with LXR ligands for 48 hours, then serum starved (1% FBS) for 1 hour prior to TCR-stimulation in PBS for 2 – 10 minutes.

#### Functional assays

To activate the TCR, cells were stimulated with 1 µg/mL plate bound anti-CD3 (UCHT1, Thermo Fisher Scientific Cat# 16-0038-85, RRID:AB_468857) and 1 µg/mL anti-CD28 (CD28.2, Thermo Fisher Scientific Cat# 16-0289-81, RRID:AB_468926,) either in solution (for 72 hour cultures) or also plate bound (for stimulations <1 hour). For microscopy experiments glass-bottomed 8-well chamber slides (Ibidi) or dishes (WillCo-dish) were coated with 5 µg/mL anti-CD3 and anti-CD28 antibodies. To measure intracellular cytokine production cells were additionally stimulated with 50 ng/mL PMA (Sigma Cat# P1585) and 250 ng/mL ionomycin (Sigma Cat# I0634) for 5 hours with GolgiPlug (BDBiosciences Cat# 555029). Ionomycin dose was increased to 1 μg/mL for IL-17A production. Donors with negligible induction of cytokines or proliferation compared to unstimulated controls were excluded from analysis (n=2).

### Lipidomics

CD3^+^ T cells were sorted by FACS and plated at 5 ×10^6^/mL into 12 well plates in complete media (n=4). A total of 10-15 ×10^6^ cells were treated with DMSO (CTRL) or GW3965 (GW, 1 µM) for 36 hours and washed twice in PBS. Frozen cell pellets were shipped to Lipotype GmbH (Dresden, Germany) for mass spectrometry-based lipid analysis as described^76^ (see Supplementary Methods). Lipidomics data has been deposited at Mendeley Data: doi: 10.17632/5rzpnr7w65.1.

### RNA sequencing and analysis

CD4^+^ T cells (3 × 10^6^) were treated with GW3965 (GW, 2 µM) for 24 hours. The LXR antagonist GSK1440233 (CTRL, 1 µM) was used as a control to supress baseline endogenous LXR activity. For TCR stimulation cells were transferred to anti-CD3/28 coated plates for the last 18 hours. Total RNA was extracted using TRIzol reagent (Life technologies) followed by DNA-free™ DNA Removal Kit (Invitrogen). RNA integrity was confirmed using Agilent’s 2200 Tapestation. UCL Genomics (London, UK) performed library preparation and sequencing (see Supplementary Methods). RNA sequencing files are available at Array Express: E-MTAB-9141.

### Analysis of gene expression

Gene expression was measured by qPCR, as in^21,77^. Primers were used at a final concentration of 100 nM. Sequences are provided in Table S3.

### Flow cytometry

Flow cytometry staining was performed as previously described^30,78,79^. (See Supplementary Methods).

### Immunoblotting

Cells were lysed in RIPA buffer and immunoblotting was performed as previously described^21^. Semi-quantitative analysis was conducted using the gel analysis module in ImageJ (National Institutes of Health, USA, RRID:SCR_003070)^80,81^.

### Chromatin immunoprecipitation

15 – 20 ×10^6^ PBMCs were rested overnight in complete media at 5×10^6^/mL in 6-well plates. Cells were treated with 1 μM GW ± 100 nM LG for 2 hours, then washed in cold PBS. Samples were double crosslinked, first with 2 mM disuccinimidyl glutarate (ThermoFisher Scientific, 20593, CAS: 79642-50-5) for 30 minutes at RT, followed by 10 minutes with 1% formaldehyde (Pierce 16% methanol free, ThermoFisher Scientific). Nuclei were isolated as previously described^21^, and chromatin was sonicated for 12 cycles of 30s on and 30s off in an ultrasonic bath sonication system (Bioruptor Pico sonication device (Diagenode)). 6 µg of pre-cleared chromatin was immunoprecipitated with 2 µg/IP anti-Histone H3K27Ac (Abcam Cat#ab4729; RRID: AB_2118291), 4 µg/IP of anti-LXRα/β (provided by Knut Steffenssen, Karolinska Institute, Sweden) or 4 µg/IP anti-rabbit IgG control (Sigma-Aldrich Cat#I5006; RRID: AB_1163659). Two IPs were performed for LXR and pooled prior to DNA purification.

To identify potential LXRE sequences we used NHR scan (RRID:SCR_016975)^82^ to interrogate the UGCG gene ±20kb, as in^21,77^. Cistrome DB was used to identify LXR ChIP-seq experiments^83,84^. Primer sequences are provided in Table S4.

### Microscopy

#### Immunostaining

CD4^+^ T cells were incubated in antibody coated chamber slides for 15 minutes at 37°C, 5% CO2 to facilitate synapse formation. Medium and non-adherent cells were discarded, and wells were washed gently with PBS before fixation (4% PFA, 2% sucrose, 140 mM NaOH, pH 7.2) for 20 minutes at RT. Formaldehyde was quenched with two washes in 0.1 M ammonium chloride (Sigma-Aldrich), followed by a PBS wash. 0.2% Trition-X-100 was used to permeabilse cells for 8 minutes at RT. Samples were blocked with 5% BSA in PBS + 0.2% fish skin gelatin (Sigma-Aldrich) overnight at 4°C. Primary antibodies were added in blocking solution for 1 hour at RT, followed by addition of fluorescently conjugated secondary antibodies for 30 minutes, RT. Cells were preserved in Prolong Diamond mounting media with DAPI (Invitrogen). For Fig. S4E fixed synapses were stained with phalloidin-FITC conjugate (Sigma).

#### Confocal microscopy

Single slices were acquired on a Leica SPE2 confocal microscope with an x63 oil-immersion objective and 488 and 633 nM excitation solid state lasers, using the following settings: 1024×1024 pixels, 600 Hz and line average of 3.

#### Total Internal Reflection Fluorescence (TIRF) Microscopy

To record live cells stained with ANE, a customised two-channel set up was used as described by Ashdown et al.^85^ and in the Supplementary Methods. 30-minute movies were acquired at a rate of 1 frame/minute. The background MFI was based on three measurements taken from the area surrounding each cell.

#### Image analysis

Image analysis was performed using ImageJ 1.51 (National Institutes of Health, USA, RRID:SCR_003070)^81^. Fluorescence intensity was analysed using the ‘Analyse Particles’ function. Mean fluorescence intensity (MFI) was measured as mean grey scale value (between 0 and 255), and corrected total cell fluorescence (CTCF) was calculated as follows: CTCF = integrated density – (cell area x MFI of background)^86^. To analyse TIRF movies of ANE-stained cells ordered and disordered channels were aligned using the Cairn Image Splitter plugin. Membrane lipid order was calculated as a GP ratio, using the plugin provided by Owen et al.^87^ at https://github.com/quokka79/GPcalc (GitHub, RRID:SCR_002630). Hue, saturation and brightness (HSB) images were set to visualize GP and pseudocoloured using the Rainbow RGB look up table.

## Statistical analysis

Statistical tests were performed in GraphPad Prism 8 (GraphPad Software, La Jolla California USA, RRID:SCR_002798, www.graphpad.com) unless otherwise stated. The D’Agostino & Pearson omnibus K2 test was used to check whether datasets were normally distributed. In some cases extreme outliers were removed based upon a ROUT test (Q=1%). Un-paired two-tailed t-tests or Mann-Whitney U were used to compare between independent groups and are represented as bar charts (mean ± SD) or violin plots (median and interquartile range). In line with previous studies on LXR agonism in human cells^9,56,88^, paired two-tailed t-tests or repeated measures ANOVA were used where cells from the same donor sample were exposed to different treatments (e.g. GW vs CTRL). This minimizes the impact of donor-to-donor heterogeneity at baseline. Where paired tests were applied, data is presented as paired line graphs. Correction for multiple comparisons was made with Tukey’s post-hoc test or Dunnet’s test (to compare all samples to vehicle), as specified. For Figure 5b p-values from multiple un-paired t-test were corrected using the two-stage linear step-up procedure of Benjamini, Krieger and Yekutieli with FDR threshold of 5%.

## Author Contributions

KEW performed most experiments and data analysis, and prepared the figures. KEW, GAR, K-SP, LMG, EC-A and SA performed flow cytometry experiments and qPCR analysis. BR-C performed western blotting experiments. DMO and II acquired microscopy images for di-4-ANEPPDHQ experiments and provided guidance for image analysis. KEW, IPT and ECJ designed experiments, interpreted the data, and prepared the manuscript. IPT and ECJ conceived the study, secured the funding and supervised all aspects of the work.

## Acknowledgments

We are grateful to K.R. Steffenson for the provision of the LXR antibody used for chromatin immunoprecipitation and to A. Castrillo and J. Thorne for earlier discussions on LXRE identification in the UGCG gene.

KEW was funded by a British Heart Foundation PhD Studentship (FS/13/59/30649) and MS Society (Grant 76). IPT was funded by a Medical Research Council New Investigator Grant (G0801278), a British Heart Foundation Project Grant (PG/13/10/30000) and an Academy of Medical Sciences Newton Advanced Fellowship. ECJ was funded by the Innovative Medicines Initiative Joint grant agreement n° 115303, as part of the ABIRISK consortium (Anti-Biopharmaceutical Immunization: Prediction and analysis of clinical relevance to minimize the risk), Arthritis Research UK Fellowships (20085 and 18106), Lupus UK, The Rosetrees Trust (M409), and University College London Hospital Clinical Research and Development Committee project grant (GCT/2008/EJ and Fast Track grant F193).

