## Supplemental Information for "LXR alters CD4^+^ T cell function through direct regulation of glycosphingolipid synthesis"

**Supplementary Information**

**Supplementary Methods**

**Lipid extraction for mass spectrometry lipidomics.** Lipids were extracted using a two-step chloroform/methanol procedure[1]. Samples were spiked with internal lipid standard mixture containing: cardiolipin 16:1/15:0/15:0/15:0 (CL), ceramide 18:1;2/17:0 (Cer), diacylglycerol 17:0/17:0 (DAG), hexosylceramide 18:1;2/12:0 (HexCer), lyso-phosphatidate 17:0 (LPA), lyso-phosphatidylcholine 12:0 (LPC), lyso-phosphatidylethanolamine 17:1 (LPE), lyso-phosphatidylglycerol 17:1 (LPG), lyso-phosphatidylinositol 17:1 (LPI), lyso-phosphatidylserine 17:1 (LPS), phosphatidate 17:0/17:0 (PA), phosphatidylcholine 17:0/17:0 (PC), phosphatidylethanolamine 17:0/17:0 (PE), phosphatidylglycerol 17:0/17:0 (PG), phosphatidylinositol 16:0/16:0 (PI), phosphatidylserine 17:0/17:0 (PS), cholesterol ester 20:0 (CE), sphingomyelin 18:1;2/12:0;0 (SM), triacylglycerol 17:0/17:0/17:0 (TAG). After extraction, the organic phase was transferred to an infusion plate and dried in a speed vacuum concentrator. 1st step dry extract was re-suspended in 7.5 mM ammonium acetate in chloroform/methanol/propanol (1:2:4, V:V:V) and 2nd step dry extract in 33% ethanol solution of methylamine in chloroform/methanol (0.003:5:1; V:V:V). All liquid handling steps were performed using Hamilton Robotics STARlet robotic platform with the Anti Droplet Control feature for organic solvents pipetting.

**Mass spectroscopy data acquisition.** Samples were analyzed by direct infusion on a QExactive mass spectrometer (Thermo Scientific) equipped with a TriVersa NanoMate ion source (Advion Biosciences). Samples were analyzed in both positive and negative ion modes with a resolution of Rm/z=200=280000 for MS and Rm/z=200=17500 for MSMS experiments, in a single acquisition. MSMS was triggered by an inclusion list encompassing corresponding MS mass ranges scanned in 1 Da increments[2]. Both MS and MSMS data were combined to monitor CE, DAG and TAG ions as ammonium adducts; PC, PC O-, as acetate adducts; and CL, PA, PE, PE O-, PG, PI and PS as deprotonated anions. MS only was used to monitor LPA, LPE, LPE O-, LPI and LPS as deprotonated anions; Cer, HexCer, SM, LPC and LPC O- as acetate adducts.

**Lipidomics data analysis.** Data were analyzed with in-house developed lipid identification software based on LipidXplorer[3,4]. Data post-processing and normalization were performed using an in-house developed data management system. Only lipid identifications with a signal-to-noise ratio >5, and a signal intensity 5-fold higher than in corresponding blank samples were considered for further data analysis. Prior to statistical analysis lipids quantified in less than 3 out of 4 donors were removed, leaving 366 lipid species. The amounts in pmoles of individual lipid molecules (species of subspecies) of a given lipid class were summed to yield the total amount of the lipid class. The amounts of the lipid classes may be normalized to the total lipid amount yielding mol%

per total lipids. Both pmol and mol% data were compared using paired t-tests. Fold changes represent the ratio of the mean of each treatment group.

**RNA sequencing.** Samples were processed using the NEB RNA Ultra II Directional assay with PolyA mRNA workflow (p/n E7760) according to manufacturer's instructions. Briefly, mRNA was isolated from 100ng total RNA using Oligo dT beads to pull down poly-adenylated transcripts. The purified mRNA was fragmented using chemical hydrolysis (heat and divalent metal cation) and primed with random hexamers. Strand-specific first strand cDNA was generated and "A-tailed" at the 3' end. Full length xGen adaptors (IDT), containing unique 8bp dual sample specific indexes, a unique molecular identifier and a T overhang are ligated to the A-Tailed cDNA. Successfully ligated cDNA molecules were then enriched with limited cycle PCR (13 cycles). Libraries to be multiplexed in the same run are pooled in equimolar quantities, calculated from Qubit and Bioanalyser fragment analysis. Samples were sequenced on the NextSeq 500 instrument (Illumina, San Diego, US) using a 43bp paired end run.

**RNA sequencing data analysis.** Run data were demultiplexed and converted to fastq files using Illumina's bcl2fastq Conversion Software v2.19. Samples were grouped by treatment or disease status. To establish differences in gene expression between groups, sequence reads were aligned using STAR v2.5.0b to the human hg19 reference genome. Gene count abundance was quantified using Partek E/M Annotation Model with default settings in Partek's RNA Flow software as in [5]. Differential expression analysis was performed using DESeq2 option in RNA Flow, which uses the Benjamin-Hochberg method for multiple testing correction. Pathway enrichment analysis was performed using Metascape [http://metascape.org][6]. Clustered heatmaps were generated in ClustVis [http://biit.cs.ut.ee/clustvis/][7], using correlation distance and average linkage for hierarchical clustering. Venn diagrams were generated with BioVenn [http://www.biovenn.nl/index.php][8].

**Flow cytometry.** 1x$10^6$ PBMCs were stained with Zombie (BioLegend) or LIVE/DEAD (ThermoFisher Scientific) fixable viability dyes for 30 minutes at 4°C, then labelled with antibodies to surface markers in cell staining buffer (PBS, 1% FBS and 0.01% sodium azide) or Brilliant Stain buffer (BD Biosciences) for 30 minutes at 4°C. Plasma membrane lipids were analysed by flow cytometry, as previously described[9]. All samples were acquired on BD LSR II or BD LSRFortessa X-20 cytometers using BD FACSDiva software. Compensation was performed using anti-mouse IgGκ/negative control compensation particles set (BD Biosciences) or OneComp eBeads (ThermoFisher Scientific), with the exception of viability dyes and filipin which were performed with single stained and unstained cells. Data was analysed using FlowJo (Tree Star). Geometric mean fluorescent intensities (gMFI) and/or percentages were exported for statistical analysis.

**Total Internal Reflection Fluorescence (TIRF) microscopy.** $CD4^+$ T cells were stained with 5 μM ANE at $1.5x10^6$ cells/mL in Hank's buffered saline solution (HBSS) with 20 μM HEPES for 30 minutes at 37°C. A Nikon Ti-E wide field microscope was used with a 1.49 NA x60 Apo-TIRF oil immersion Nikon objective, under TIRF conditions and an Andor sCMOS camera for signal capture. To record live cells stained with ANE a customised two-channel set up was used to simultaneously record signal from ordered and disordered membranes. 30-minute movies were acquired at a rate of 1 frame per minute. A 488 nm laser set to 40% power was used for the excitation. Emitted light was directed to a Cairn OptiSplit III 2-channel image splitter, equipped with a 605 nm dichroic mirror,and 542/50 nM and 660/52 nM bandpass filters. This separated the fluorescent signal into ordered (542/50) and disordered (660/52) channels prior to detection. A suitable correction lens and neutral density filter were used to ensure similar intensity in both channels.

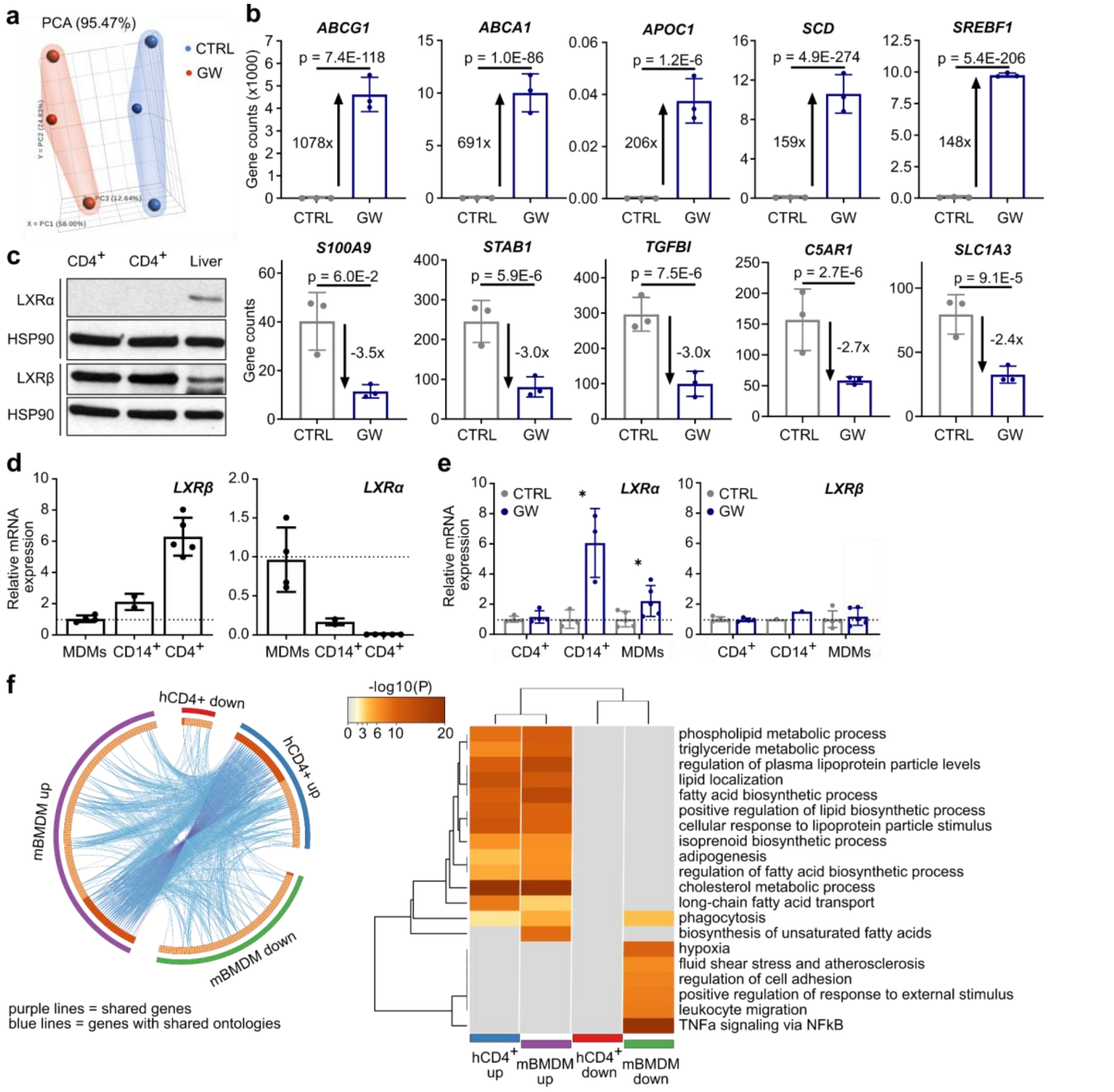


**Fig. S1. (a)** Principal component analysis (PCA) comparing RNA-seq data from CD4$^+$ T cells treated with LXR agonist GW3965 (GW) for 24 hours to control (CTRL). **(b)** Normalised RNAseq gene counts for the genes with the greatest response to GW stimulation. **(c)** LXRα and LXRβ protein expression by Western blotting in CD4+ T cells (n=2) and human liver lysate with HSP90 loading control. **(d)** mRNA expression of LXRα and LXRβ was measured in cells from independent donors: human monocyte derived macrophages (MDMs) (n=4), CD14$^+$ monocytes (n=2) or CD4$^+$ T cells (n=5). Gene expression was normalized to cyclophilin A and expressed relative to MDMs (MDMs=1, shown by dashed line). **(e)** Human immune cells were cultured with 1 μM GW for 16-18 hours (n=3-9). Expression of LXR subtypes was analysed by qPCR (CTRL=1, shown by dashed line). **(f)** Comparison of GW-regulated genes in human CD4$^+$ T cells and murine macrophages. The circos plot shows GW-regulated genes (purple lines) and functional pathways (blue lines) common to human CD4$^+$ T cells and murine bone marrow derived macrophages (BMDMs)(5). The clustered heatmap compares the statistical significance of the top 20 enriched pathways across the two cell types. **(b-e)** All histograms show mean ± SD. Unpaired two-tailed t-tests; $*p < 0.05$.

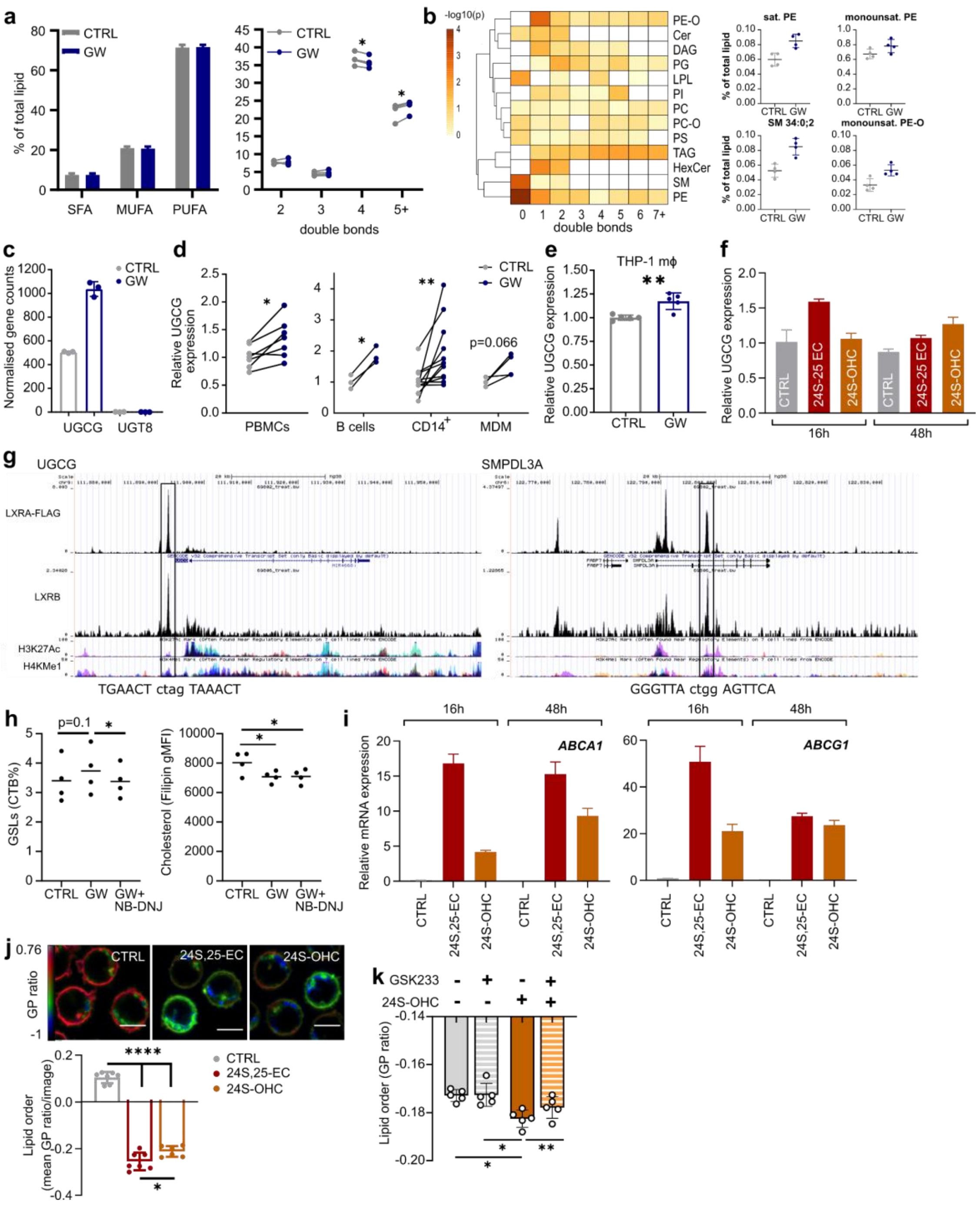


**Fig. S2. (a-b)** Lipid saturation was measured by shotgun lipidomics. Percentage of lipids with each degree of unsaturation was compared in total **(a)** and within each lipid class **(b)**. Heatmap shows p-values of comparisons made within subclasses with dot plots of the top hits. White squares represent absent values. **(c)** Normalised RNA-Seq gene counts for UGCG and UGT8. **(d-f)** UGCG mRNA expression was measured by qPCR. **(d)** PBMCs (n=7, 5 hours), B cells (n=3), CD14+ monocytes (n=12) and monocyte-derived macrophages (MDM) (n=4), **(e)** differentiated THP-1 macrophages were stimulated with GW and **(f)** T cells treated with endogenous LXR ligands 24S, 25-epoxycholesterol (24S-25 EC) or 24S-hydroxycholesterol (24S-OHC) compared to ethanol control (CTRL). **(g)** LXR occupancy in HT29 colorectal cancer cells treated with GW for 2 hours (10 μM) [10] and ENCODE data of histone marks commonly associated with regulatory sequences. Highlighted peaks labelled with putative DR4 sequences identified with NHR scan show regions amplified by ChIP-qPCR. **(h)** Cholesterol and glycosphingolipid levels measured in cell treated with GW ± UGCG inhibitor (NB-DNJ) for 24 hours (n=4). **(i)** Induction of LXR target genes ABCA1 and ABCG1 was analysed by qPCR (n=3, pooled). **(j)** Membrane lipid order was analysed with di-4-ANEPPDHQ (n=1) and the mean GP ratio per image was quantified and compared. Scale bar represents 5 μM. **(k)** Cumulative data from three experiments showing lipid order measured by flow cytometry. Cells were treated with an LXR agonist (24S-OHC) ± LXR antagonist (GSK233)(n=5) for 24 hours. Data is shown as mean ± SD. **(a-e, h)** Two-tailed t-tests and **(j-k)** One-way ANOVA with Tukey's posthoc test *p < 0.05, **p < 0.01, ****p < 0.0001.

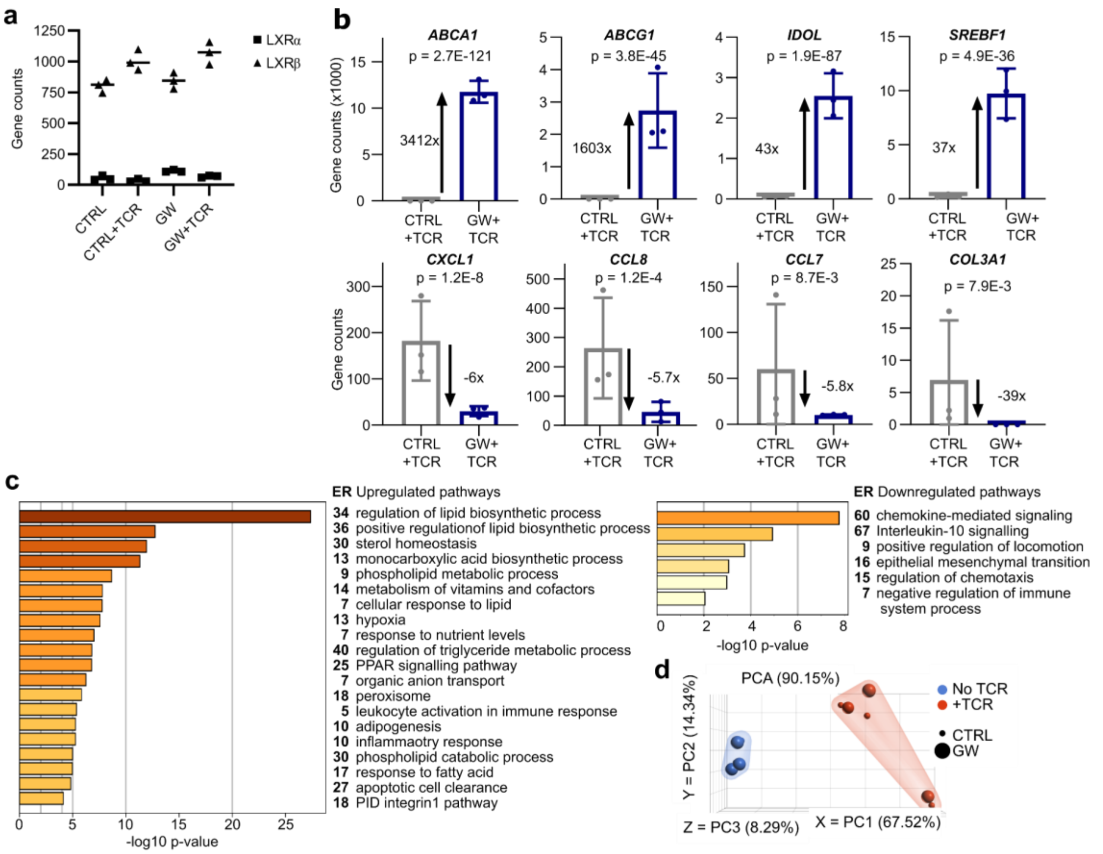


**Fig. S3.** RNA-seq was performed on CD4+ T cells (n=3) incubated ± GW for 6 hours, before stimulation with anti-CD3/CD28 (TCR) ± GW. **(a)** Normalised gene counts comparing the expression of LXRα and LXRβ and **(b)** of differentially expressed genes showing the strongest differences. **(c)** Pathway enrichment analysis of genes up- or down-regulated in GW+TCR compared to CTRL+TCR. Bar chart plots p-values and is annotated with the enrichment ratio (ER). **(d)** Principal component analysis (PCA) comparing TCR stimulated to unstimulated samples.

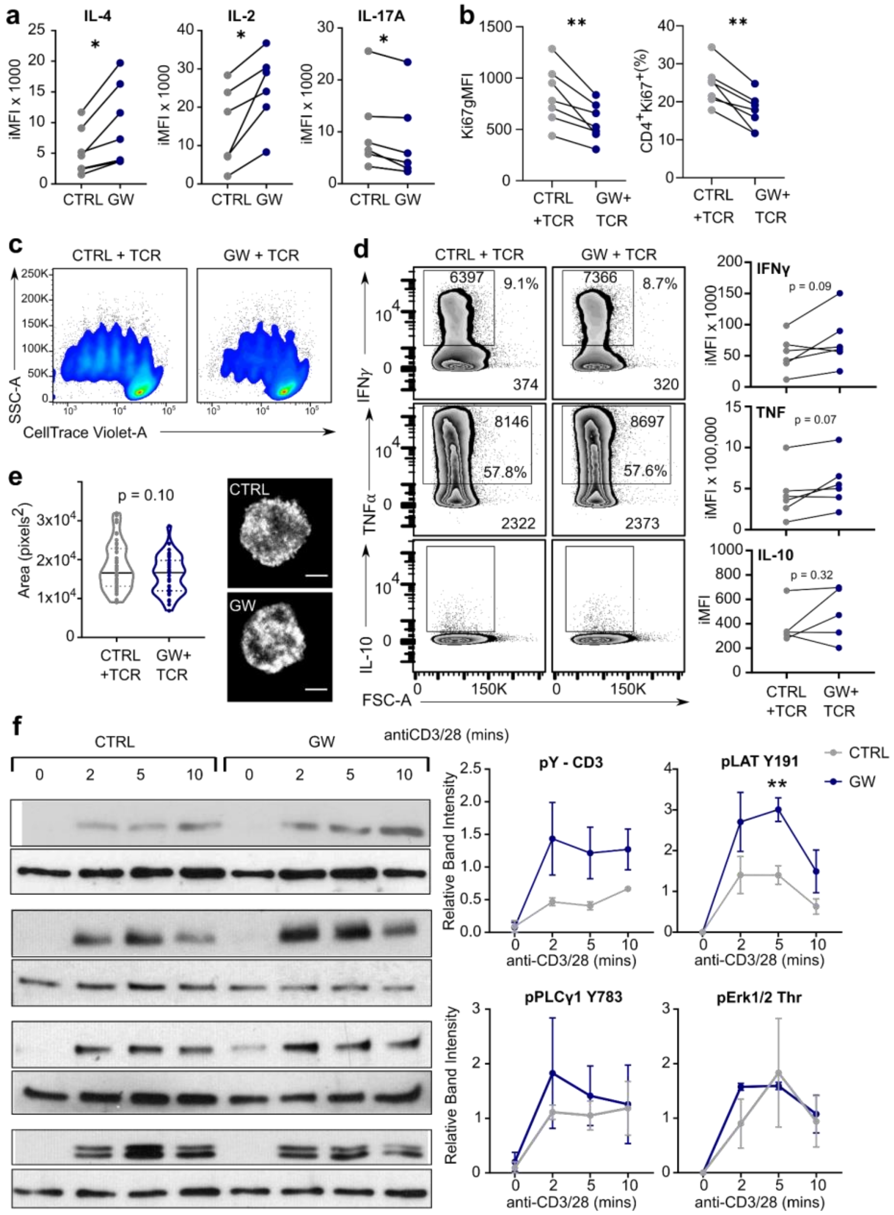


**Fig. S4.** CD4⁺ T cells were activated with anti-CD3/CD28 (+TCR) for 72 hours in the presence of GW3965 (GW) or control (CTRL). Intracellular cytokines were analysed by flow cytometry after additional treatment with PMA and ionomycin. **(a)** Cumulative data from four independent experiments shows cytokine production expressed as an integrated MFI (iMFI = gMFI*frequency of cytokine producing cells [11]). **(b-c)** Proliferation was measured using Ki67 **(b)** and dilution of CellTraceViolet **(c)**. Cumulative data from four independent experiments **(b)**, and flow cytometry plot representative of n=3 **(c)**. **(d)** Flow cytometry plots labelled with percentage of positive cells, and the geometric mean fluorescence intensities (gMFI) of both the cytokine-producing population (inside gate) and whole population (bottom left) are shown. Cumulative data from 4 independent experiments. **(e)** $CD4^+$ T cells were activated with antibody coated coverslips for 10 minutes. Fixed synapses were stained with phalloidin and imaged by TIRF microscopy to assess the area of the cell-coverslip interface (n >16 cells/donor, n= 2 donors). **(f)** Signaling protein phosphorylation after 2, 5 and 10 minutes of TCR engagement by Western blotting. Band intensities relative to HSP-90 (as loading control) were calculated for 2-3 independent experiments (n=2-4 donors). Two-tailed t-tests: $^*p<0.05$, $^{**}p<0.01$. Abbreviations: Lck – lymphocyte-specific protein tyrosine kinase; pY – phosphotyrosine; CD3 – cluster of differentiation 3; LAT – linker for activation of T cells; PLCγ – phospholipase Cγ1; Erk – extracellular signal related kinase; HSP90 – heat shock protein 90.

**Table S1. Identification of novel LXR-regulated transcripts.** To determine whether genes had previously been linked to LXR a literature search conducted in PubMed using terms ‘LXR’, ‘NR1H2’, ‘NR1H3’ and gene symbols. The T-cell only gene list was also cross-referenced with published data from human macrophages[12], THP-1 cells[13] and BMDMs[14].

| | |
|---|---|
| No direct link to LXR identified | BRWD3, CHD2, MKNK2, SLC29A2, TDRD6, TGFBI, TKT, UGCG |
| Evidence of LXR regulation | IFI30, CD14 and C5AR1[14], S100A9[15], AQP9[16], MSMO1[17], ABCD1[13], MMAB[12], ACSL1[18] |
| Evidence of species-specific LXR regulation | SMPDL3A[13], OLMALINC[19], IL1RN[12], TGFBI[12] |

**Table S2. Interaction of LXR and TCR activation on gene expression.** LXR-regulated genes were hierarchically clustered, and four distinct patterns of gene expression were identified based on their response to LXR activation with GW3965 and T cell receptor stimulation (anti-CD3/28). For each cluster (A-D) the responses to LXR ligand (LXR) and TCR stimulation (TCR) are summarized and all genes within the cluster are listed.

| | |
|---|---|
| Cluster A<br>LXR: up<br>TCR: none | RDH11, SLC25A1, STX1A, LILRA1, LINC01578, C3, STARD4, CLCN6, MYLIP, SREBF1, ABCA1, INSIG1, ABCD1, ACSL3, TMEM135, ABCG1, MID1IP1, SCD, BRWD3, OLMALINC, RARA, FADS1, FADS2 |
| Cluster B<br>LXR: up<br>TCR: down | APOE, SMPDL3A, LSS, MKNK2, NR1H3, SLC29A2, EEPD1, CD14, TLR4, FBP1, IDH1, FGR, KMO, SDC2, ALDOC, GAPT, CORO7, LOC100130872, BLOC1S1-RDH5, LEF1, ZNF775 |
| Cluster C<br>LXR: up<br>TCR: up | ISY1-RAB43, TNFSF15, PLAUR, TGM2, ADM, UGCG ATP5J2-PTCD1, METTL9, SREBF2, LDLR, LPCAT3, MVD, RNF145, GPR82, PCYT2, PRDX5, ACACA, DBI DHCR7, FASN, MMAB, MVK, FDPS, TNC, IL1RN, TCF15, TMEM160, LIPG, ZBTB10, FAM213A, OLR1 |
| Cluster D<br>LXR: down/none<br>TCR: up/down | SLFN12L, SUCNR1, ACOD1, LNPEP, MSH5, PHOSPHO2-KLHL23, CCL2, CCL7, CCL8, COL3A1, PCDH7, TENM4, DOCK4, CXCL1, CXCL2, LRRC24, RPS10-NUDT3, GNRHR, ZNF66, LOC100507053, SAMD12, CORO7-PAM16, LOC101593348, TGFBI, TNS3, MIAT, TRIM39-RPP21 |

**Table S3.** Oligonucleotide sequences for qPCR.

| Gene | Forward primer (5'-3') | Reverse primer (5'-3') |
|---|---|---|
| ABCA1 | TGAGCTACCCACCCTATGAACA | CCCCTGAACCCAAGGAAGTG |
| ABCG1 | TGCAATCTTGTGCCATATTTGA | TGCAATCTTGTGCCATATTTGA |
| Cyclophilin A (*PPIA*) | GCATACGGGTCCTGGCATCTTGTCC | ATGGTGATCTTCTTGCTGGTCTTGC |
| FASN | CTGCTGCTGGAAGTCACCTA | GTGTGTGTTCCTCGGAGTGA |
| LXRα (*NR1H3*) | AGAGGAGGAACAGGCTCATG | AAAGGAGCGCCGGTTACACT |
| LXRβ (*NR1H2*) | GGAGCTGGCCATCATCTCA | GTCTCTAGCAGCATGATCTCGGATAGT |
| OLMALINC* | GACTCCTTTGG GAGACCAGTG | AGGTCACAGGGGATTTGATGG |
| SCD | GCAAACACCCAGCTGTCAAA | GCACATCATCAGCAAGCCAG |
| SREBP1c (*SREBF1*) | TCAGCGAGGCGGCTTTGGAG | CATGTCTTCGATGTCGGTCAG |
| UGCG | CGTCCTCTTCTTGGTGCTGT | AGAGAGACACCTGGGAGCTT |

*Oligonucleotide sequences from[20].

**Table S4.** Oligonucleotide sequence for ChIP-qPCR.

| Target | Forward primer (5'-3') | Reverse primer (5'-3') |
|---|---|---|
| UGCG DR4 | ACTCTAGTCACTCCCCTGGAC | GCCTGATCTTGATAAACCACTGG |
| SMPDL3A LXRE* | TGCAATCTTGTGCCATATTTGA | TGCAATCTTGTGCCATATTTGA |

*Oligonucleotide sequences from[13].

**Dataset S1 (separate file).** Differentially expressed genes and Metascape pathway analysis results for GW vs GSK233 comparison.

**Dataset S2 (separate file).** Differentially expressed genes and Metascape pathway analysis results for GW+TCR vs GSK233+TCR comparison.
